# Basal forebrain cholinergic circuits orchestrate diverse cell types in the adult dentate gyrus to support neural stem cell function and spatial memory

**DOI:** 10.1101/2022.05.25.493227

**Authors:** Luis Quintanilla, Yijing Su, Jeremy M. Simon, Yan-Jia Luo, Brent Asrican, Seth Tart, Ryan N. Sheehy, Ya-Dong Li, Guo-li Ming, Hongjun Song, Juan Song

**Author notes:** Correspondence should be addressed to: Juan Song, Ph.D.

## Abstract

Dentate gyrus (DG) is a critical structure involved in spatial memory and adult neurogenesis, two distinct processes dynamically regulated by local circuits comprising diverse populations of DG cells. It remains unknown how these DG cells are orchestrated to regulate these distinct hippocampal functions. Here we report activation of a cholinergic circuit from the Diagonal Band of Broca to DG promotes quiescent radial neural stem cell (rNSC) activation and spatial memory. Furthermore, single-nucleus RNA-sequencing reveals broad transcriptomic changes across DG mature and adult-born cells in response to cholinergic-circuit activation. Notably, neuronal populations exhibit cholinergic-activity-induced molecular changes related to synaptic functions crucial for spatial memory; while rNSCs exhibit changes related to structural remodeling and neurogenic proliferation crucial for hippocampal neurogenesis. Electrophysiology and NicheNet analyses reveal granule cells, endothelial cells, and astrocytes as potential intermediaries for cholinergic regulation of rNSCs. Our findings reveal cell-type-specific signaling mechanisms underlying cholinergic regulation of distinct DG functions.

## Introduction

The dentate gyrus (DG) is a unique brain structure that produces newborn neurons in adulthood and participates in spatial memory. As a neurogenic region, adult DG contains endogenous quiescent radial neural stem cells (rNSCs) that provide a self-renewable source for the continuous replenishment of new neurons throughout life. Adult-born neurons derived from rNSCs make functional contributions to memory and emotion processing (Anacker et al., 2018; Sahay et al., 2011; Santarelli et al., 2003; Snyder et al., 2005). As a center for learning and memory, the DG is the first input region in the hippocampal tri-synaptic circuit, in which DG granule cells (GCs) receive major excitatory inputs from the entorhinal cortex (EC) and relay information to CA3 through mossy fibers (Bao and Song, 2018; Knierim and Neunuebel, 2016; Treves et al., 2008; Vivar and van Praag, 2013). The sparse firing properties of GCs are critical for processing spatial information and pattern separation (Knierim and Neunuebel, 2016). Importantly, both learning/memory and adult neurogenesis processes are dynamically regulated by neural circuits comprising diverse populations of DG cells. The major neuronal types in the DG include GCs, interneurons, and mossy cells, which all have been shown to play critical roles in certain forms of learning and memory, including spatial memory and pattern separation (Andrews-Zwilling et al., 2012; Bui et al., 2018; Deng et al., 2009; Favuzzi et al., 2017; Li et al., 2021; Madronal et al., 2016). Moreover, these cells also serve as niche cells for rNSCs to control the neurogenesis process. We recently showed that DG parvalbumin (PV) and cholecystokinin (CCK) interneurons and hilar mossy cells can regulate the activation/quiescence state of rNSCs and subsequent production of newborn progeny (Asrican et al., 2020; Song et al., 2012; Yeh et al., 2018). Besides neuronal cells, glia or endothelial cells have also been shown to be regulated by local DG interneurons, which in turn influences rNSC behavior and neurogenesis (Asrican *et al*., 2020; Shen et al., 2019). How diverse populations of DG cells are orchestrated to regulate distinct hippocampal functions at the circuit and molecular levels remains elusive.

To address this question, we focused on a basal forebrain cholinergic circuit based on its reported broad actions on multiple DG cell types (Pabst et al., 2016). We hypothesize that the basal forebrain cholinergic circuit may recruit both neuronal and non-neuronal components in the DG to differentially regulate spatial memory and adult neurogenesis. The source of acetylcholine (Ach) in the DG originates from the Medial Septum (MS) / Diagonal Band of Broca (DB) complex within the basal forebrain and is thought to play an important role in hippocampal dependent learning and memory (Deiana et al., 2011; Hasselmo, 2006). Interestingly, our anterograde projection tracing showed that the DB sent more projections to DG than MS. Importantly, optogenetic stimulation of DB-DG (but not MS-DG) cholinergic projections promotes activation of rNSCs. Moreover, optogenetic activation of the DB-DG cholinergic projections during spatial memory encoding promotes spatial memory retrieval. To further address how DB cholinergic inputs regulate distinct cellular components, we performed single-nucleus RNA-seq of the DG and found broad transcriptomic changes across DG mature and adult-born cells in response to activation of the DB-DG cholinergic circuits. We identified cholinergic-activity-induced molecular changes related to synaptic functions in various mature neuron populations and those related to structural remodeling and neurogenic proliferation in rNSCs. Furthermore, we performed slice electrophysiology and NicheNet analyses to address the niche cells that potentially mediate cholinergic activity-dependent regulation of rNSCs, and identified mature GCs, endothelial cells, and astrocytes as potential intermediaries for such regulation.

## Results

### The Diagonal Band of Broca sends more cholinergic projections to the Dentate Gyrus than the Medial Septum

In the basal forebrain, cholinergic neurons are mainly located in the MS and DB (Nyakas et al., 1987; Senut et al., 1989) and these neurons send dense projections to the hippocampal formation (Wu et al., 2014). To determine whether there are differences between MS and DB cholinergic projections to the DG, we performed anterograde projection tracing of MS and DB cholinergic neurons and compared their projections in the DG. Cre-dependent AAVs expressing mCherry (to the MS) and YFP (to the DB) was injected into mice expressing Cre under the promoter of choline acetyltransferase (ChAT-Cre) (Rossi et al., 2011) for selective targeting of cholinergic neurons (Figure 1A, Figure S1). Reporter expression was found to be restricted to cholinergic cells expressing ChAT in the MS and DB (Figure 1B). Significantly more YFP+ ChAT+ cells were labeled in the DB than mCherry+ ChAT+ cells in MS (Figure 1E and 1F) with similar labeling efficiency (Figure 1G), suggesting that more cholinergic neurons exist in the DB as compared to MS. Interestingly, we found significantly more DB-DG cholinergic projections in the DG/hilus and the molecular layer than MS-DG projections (Figure 1D and 1H). Importantly, these DB-DG cholinergic projections from DG/molecular layer exhibited close association with rNSCs (Figure 1I and 1J). These results suggested that DB cholinergic inputs may exert more control over DG and rNSCs than MS cholinergic inputs.

**Figure 1:**
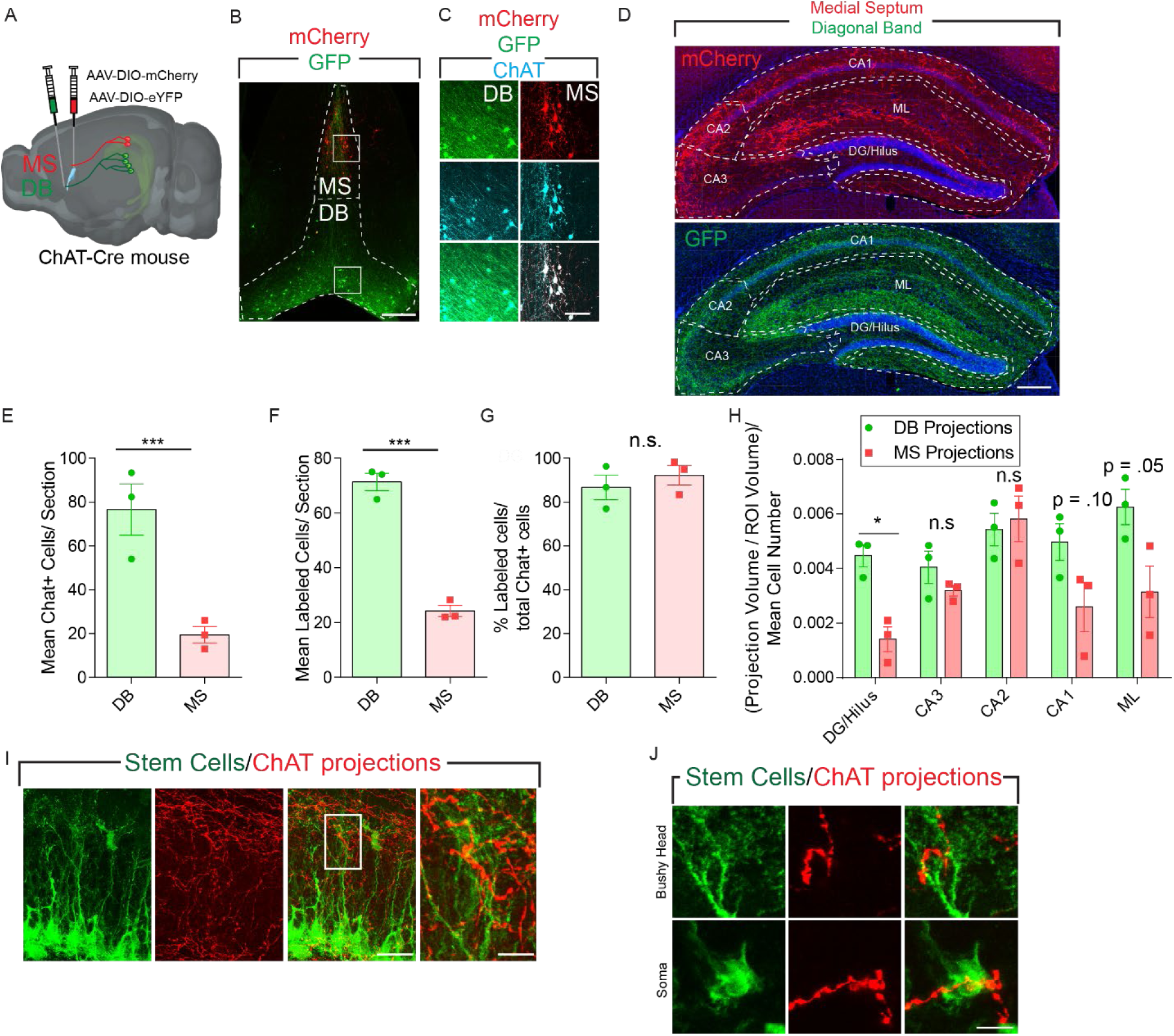
Diagonal Band of Broca send more cholinergic projections to DG than medial septum. A) Schematic representing experimental design using AAV’s to target MS and DB using distinct color fluorophores for anterograde projection quantification. B) Immunofluorescence image of AAV injection site to the MS/DB. Scale bar 200 µm C) Colocalization of ChAT+ cells with distinct reporter fluorophores in the MS and DB. Scale bar 50 µm. D) Immunofluorescence image of hippocampus with ROIs drawn around respective hippocampus subregions. ROIs depicted were used for analysis of figure 1H. Scale bar 100 µm. E) Quantification of mean ChAT+ cells in each immunofluorescence section. Bars indicate mean +/- S.E.M. p =.0097 n = 3 by Student’s t-test. F) Quantification of mean labeled cells in each section Bars indicate mean +/- S.E.M. p =.0002 n = 3 by Student’s t-test. G) Percent of labeled cells or labeling efficiency of our AAV approach. Bars indicate mean +/- S.E.M. n = 3. H) Normalized projection density to respective hippocampal regions from MS and DB. Bars indicate mean +/- S.E.M. p =.0072, q = .0366, by multiple t-tests with Bonferroni correction n = 3. I) Confocal microscopy image of cholinergic fibers associating with rNSCs. Left Scale bar = 10 µm, right scale bar = 1 µm. J) Immunofluorescence image of Nestin:GFP positive cell associating with cholinergic fibers expressing an mCherry reporter with different subdomains of an adult rNSC in the SGZ. Scale bar = 5 µm.

### Stimulation of DB-DG cholinergic circuits promotes rNSC activation

We next sought to determine the functional effects of DB and MS cholinergic projections on rNSC behavior. We first took a gain-of-function approach by optogenetic activation of DB-DG or MS-DG projections in vivo (blue light pulses, 8 Hz, 5 ms, 30 s/5 min, 8 hours/day for 3 days) by delivering Cre-dependent AAVs expressing ChR2 or YFP in DB or MS of ChAT-Cre mice (Figure 2A, 2B, 2I, and 2J). The use of 8 Hz was based on the known firing frequency of forebrain cholinergic neurons during exploration, which relates to the hippocampal theta rhythm mediated by the MS/DB (Colgin, 2016; Vanderwolf, 1969; Zhang et al., 2010). Using immunohistology and confocal microscopy, we assessed the proliferation of rNSCs by injecting the thymidine analog EdU on the last day of light stimulation to label proliferating cells. As a result, DB-DG cholinergic circuit stimulation led to a significant increase in the density of proliferating rNSCs (nestin+ EdU+) (Figure 2C - 2E) and the proliferating rate of rNSCs (Figure 2F) without significantly altering the densities of proliferating progeny (EdU+) (Figure 2G) and rNSC pool (nestin+) (Figure 2H). These results suggest that activation of the DB-DG cholinergic circuit promotes the proliferation of rNSCs. Additionally, we validated these effects with a different approach by chemogenetic activation of DB cholinergic neurons in vivo. Specifically, we injected Cre-dependent AAVs expressing excitatory DREADDs hM3Dq to the DB, and CNO water was given for 3 days followed by EdU injection and proliferation assay (Figure S2A and S2B). Consistent with the results obtained using optogenetics, we observed a significant increase in the proliferation of rNSCs upon chemogenetic activation of DB cholinergic neurons (Figure S2C – S2H). By contrast, optogenetic activation of MS-DG cholinergic projections with a similar stimulation paradigm did not alter rNSC proliferation (Figure 2I - 2P). These results support the unique contribution of DB-DG cholinergic circuits in regulating activation of rNSCs.

**Figure 2:**
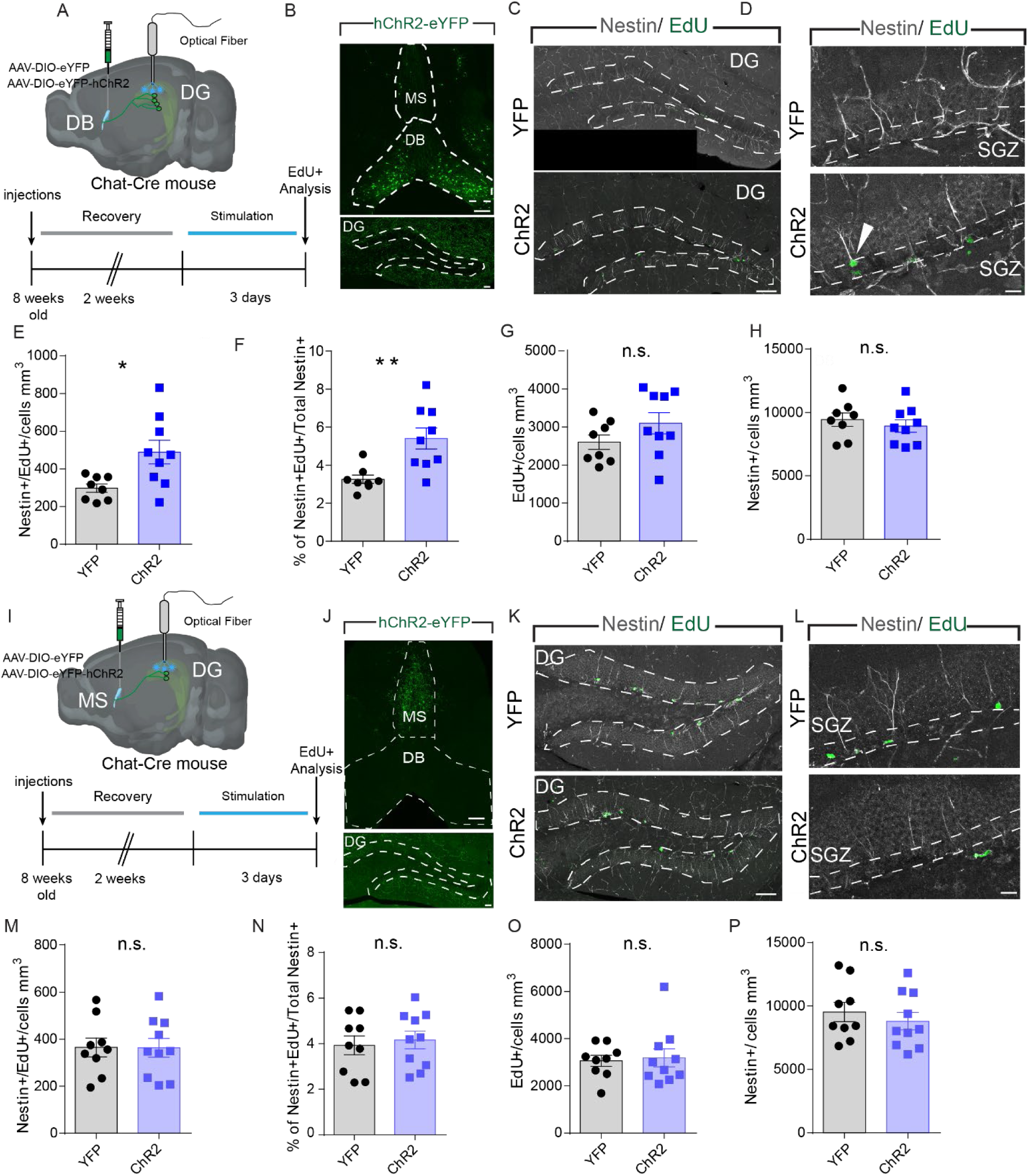
Stimulation of DB-DG (but not MS-DG) cholinergic circuits promotes rNSC activation. A) Schematic of timeline and experimental setup for optogenetic stimulation of DB cholinergic afferents and rNSC proliferation analysis. B) Immunofluorescence image of DB targeted AAV viral injection and ChR2 afferents in the DG. DB scale bar, DG Scale bar 100 µm 20 µm. C) Representative immunofluorescence section of DG used to quantify rNSC proliferation with Nestin/EdU. Scale bar = 100 µm D) Representative immunofluorescence section of SGZ used to quantify rNSC proliferation. Scale bar = 10 µm E) Quantification of proliferating neural stem cells. Bars indicate mean +/- S.E.M. n= (8,9), p-val = .0155 by student’s t-test . F) Quantification of percent of proliferating neural stem cells. Bars indicate mean +/- S.E.M. n= (8,9), p-val = .0037 by student’s t-test . Quantification of neural stem cell pool. Bars indicate mean +/- S.E.M n = (8,9). G) Quantification of overall proliferation in the SGZ. Bars indicate mean +/- S.E.M. n = (8,9). H) Quantification of neural stem cell pool. Bars indicate mean +/- S.E.M n = (8,9). I) Schematic of timeline and experimental setup for optogenetic stimulation of MS cholinergic afferents and rNSC proliferation analysis. J) Immunofluorescence image of MS targeted AAV viral injection and ChR2 afferents in the DG. DB scale bar 100 DG Scale bar 20 µm. K) Representative immunofluorescence section of DG used to quantify rNSC proliferation. Scale bar = 100 µm L) Representative immunofluorescence section of SGZ used to quantify rNSC proliferation with Nestin/EdU. Scale bar = 10 µm M) Quantification of proliferating neural stem cells. Bars indicate mean +/- S.E.M. n= (9,10) N) Quantification of neural stem cell pool in the SGZ. Bars indicate mean +/- S.E.M. n = (8,9). O) Quantification of overall proliferation in the SGZ. Bars indicate mean +/- S.E.M. n = (9,10). P) Quantification of percent of proliferating neural stem cells. Bars indicate mean +/- S.E.M. n= (9,10).

### Ablation of DB cholinergic neurons decreases rNSC activation

To address whether the DB cholinergic neurons are required for rNSC proliferation, we took a loss-of-function approach by selectively ablating DB cholinergic neurons. Cre-dependent AAVs expressing Caspase3 or YFP were injected into the DB of ChAT-Cre mice, and 6 weeks later, a 70% reduction of DB ChAT+ neurons were observed in Caspase-injected mice as compared to YFP-injected mice, indicative of efficient ablation of cholinergic neurons (Figure 3A - 3C). To assess the proliferation of rNSCs, we injected EdU to label proliferating cells 6 weeks after cholinergic neuron ablation. As a result, ablation of DG cholinergic neurons led to a significant reduction in the density of proliferating rNSCs (Figure 3D and 3E) and the proliferating rate of rNSCs (Figure 3F) without altering the densities of proliferating progeny (Figure 3G) and rNSC pool (Figure 3H). These results suggest that DB cholinergic neurons are required for rNSC proliferation.

**Figure 3:**
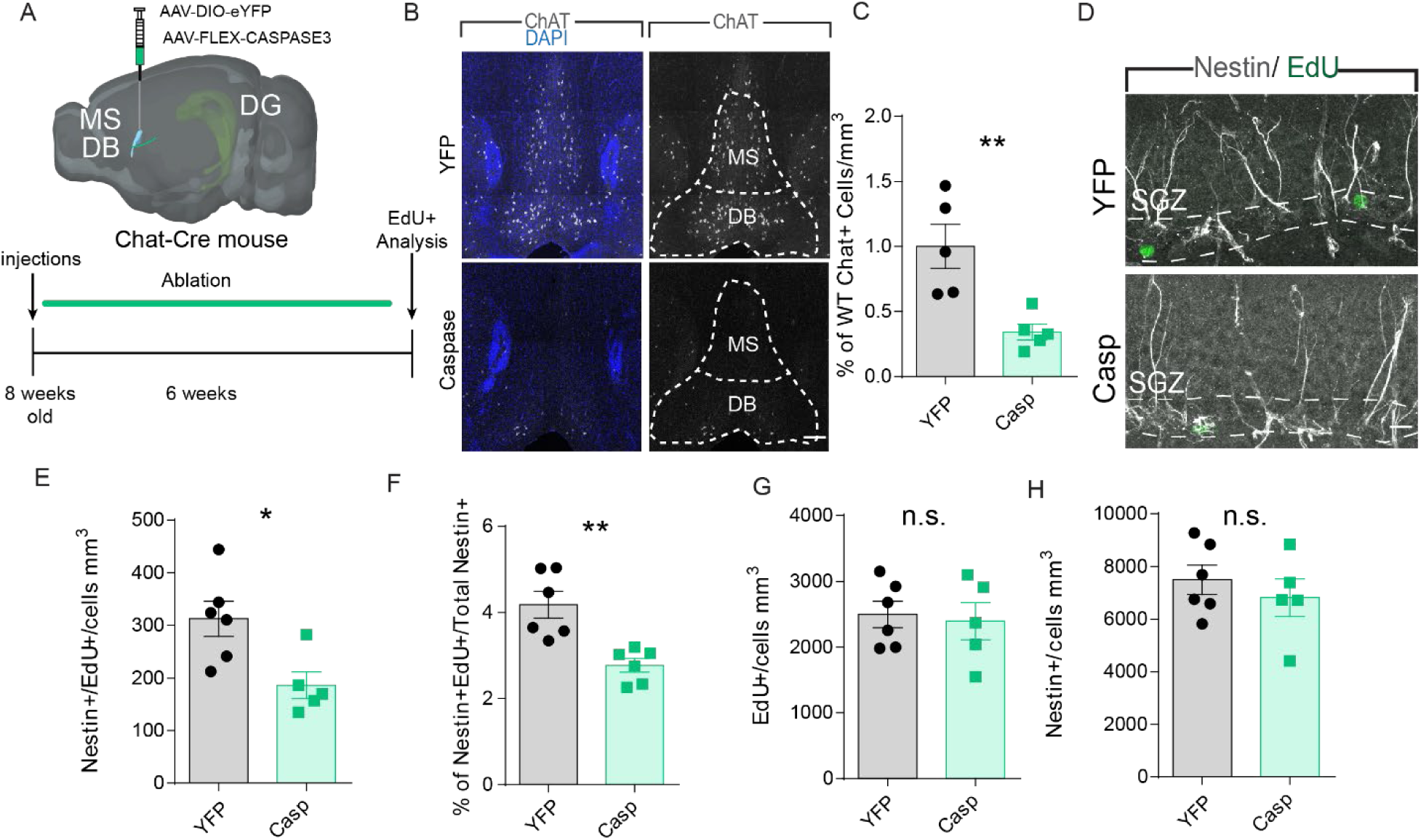
Ablation of DB cholinergic neurons decreases rNSC activation. A) Schematic representing ablation strategy used to target the MS/DB and ablate cholinergic cells. B) Representative image of MS and DB ChAT positive cells in control and caspase groups. Scale bar = 100 µm C) Quantification of percent of Mean ChAT+ positive cells compared to YFP animals. Bars indicate mean +/- S.E.M. n = 5, p-val = .0062 by student’s t-test D) Representative immunofluorescence image of Nestin+ EdU+ cells in SGZ used to quantify the effects of cholinergic ablation on rNSCs. Scale bar = 10 µm E) Quantification of proliferating neural stem cells showing decrease after MS/DB ablation. Bars indicate mean +/- S.E.M. n= (6,5) p-val =.0174 student’s t-test F) Quantification of percent of proliferating neural stem cells. Bars indicate mean +/- S.E.M. n = (6,5) p value = .0024 student’s t-test G) Quantification of overall proliferation after ablation of MS/DB. Bars indicate mean +/- S.E.M. n = (6,5) H) Quantification of neural stem cell pool after MS/DB cholinergic ablation. Bars indicate mean +/- S.E.M. n = (6,5)

### Stimulation of DB-DG cholinergic circuits during spatial memory encoding promotes spatial memory performance

Given that ACh plays an important role in memory encoding (Buccafusco et al., 2005; Green et al., 2005; Hasselmo, 2006; Winkler et al., 1995), we wondered whether the DB-DG cholinergic circuit impacts the ability of animals to encode memories. To address this question, we adopted two hippocampus-dependent memory tasks: novel place recognition (NPR) for spatial memory, and contextual fear conditioning (CFC) for contextual memory. Specifically, we injected Cre-dependent AAVs expressing ChR2 or YFP to the DB of ChAT-Cre mice and implanted optical fibers bilaterally above the DG where we stimulated DB-DG cholinergic projections at 8Hz during spatial or contextual memory encoding (Figure 4A – 4C). For the NPR test, a significant increase in the discrimination ratio and exploration time with the object associated with the novel location was observed in ChR2-injected mice (Figure 4D and 4E), suggesting improved spatial memory retrieval. Importantly, mice did not show biases for either object during light stimulation (Figure 4F). In addition, no differences were observed in time spent at the center of an open field test (Figure 4G) or locomotion during encoding or retrieval (Figure S3) when comparing ChR2 and YFP mice, indicating that ChR2 mice were not anxious, and their locomotor activity was normal. By contrast, for the CFC test, no significant differences in the freezing time at 1 hour and 24 hours after foot shocks were observed when comparing ChR2 and YFP mice (Figure 4H - 4J). Together, these results suggest that the DB-DG cholinergic circuit plays a critical role in spatial (but not contextual) memory encoding. Specifically, cholinergic circuit activation during spatial memory encoding leads to improved spatial memory performance.

**Figure 4.**
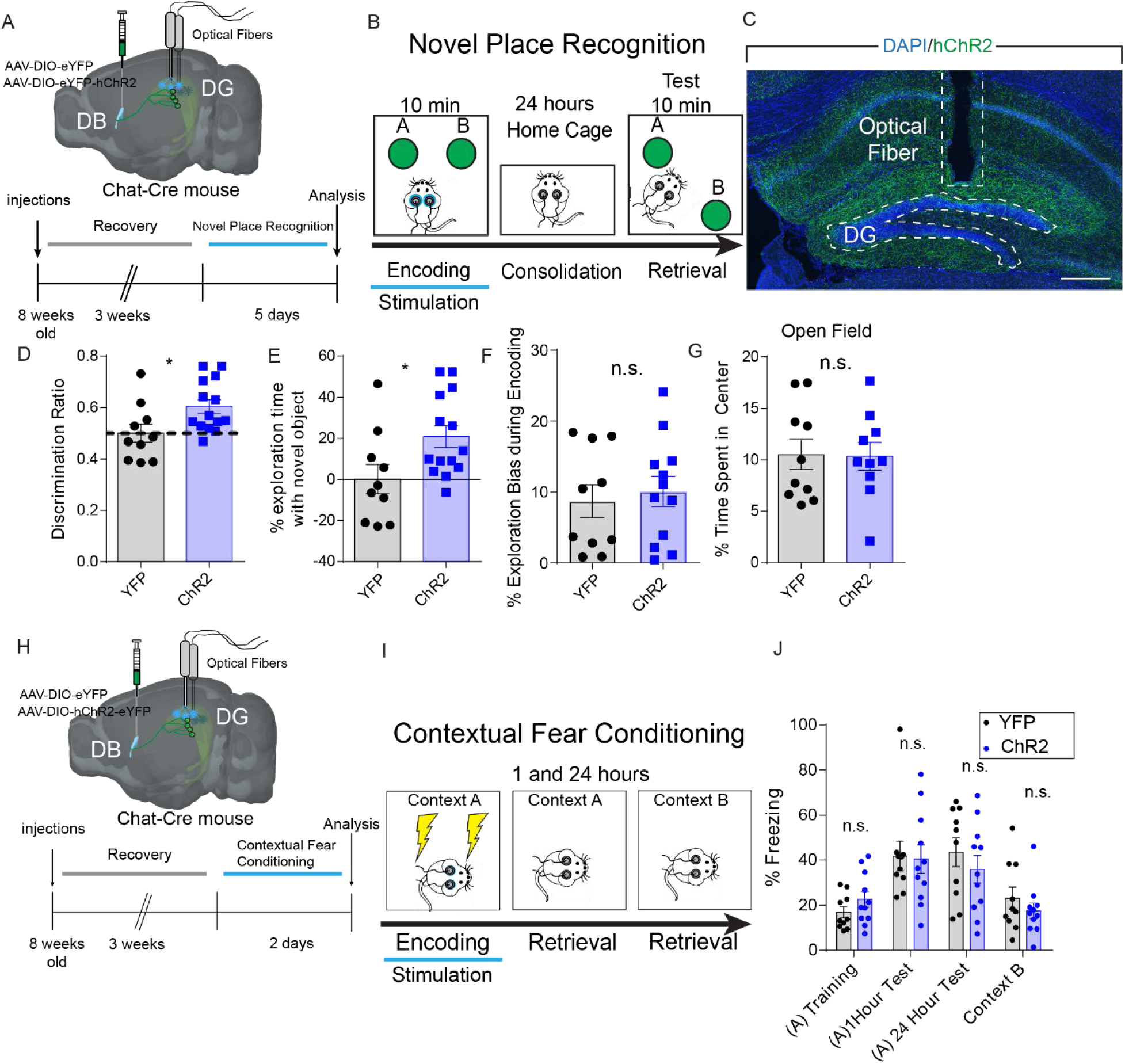
Stimulation of DB-DG cholinergic circuits during spatial memory encoding promotes spatial memory performance. A) Schematic representing behavioral paradigm used during novel place recognition. B) Novel place recognition paradigm used to test encoding of spatial memories. Animals were stimulated during the encoding phase of the task. C) Representative immunofluorescence image of optical fiber implantation site above the DG. Scale bar 100 µm. D) Quantification of discrimination ratio used to test spatial memory between old location and new location object (see methods for calculations). Bars indicate mean +/- S.E.M. n = (10,14) p-val =.0259 student’s t-test E) % Exploration time with novel location object used to test spatial memory (see methods for calculations). Bars indicate mean +/- S.E.M. n = (10,14) p-val =.0259 student’s t-test F) % Exploration Bias during encoding. Bars indicate mean +/- S.E.M. n = 10 G) Open Field test used to test anxiety during stimulation. Bars indicate mean +/- S.E.M. n = 10 H) Schematic for contextual fear conditioning paradigm used to evaluate primarily non-spatial memory. I) Contextual fear conditioning paradigm used to evaluate short-term and long-term memory encoding by stimulating DB afferents during memory acquisition. J) Quantification of % freezing during all aspects of contextual memory task. Bars indicate mean +/- S.E.M. n = 10

### Single-nucleus RNA-seq reveals DG heterogeneity

Our results showed that stimulating the DB-DG cholinergic circuit is critical for regulating two distinct DG-dependent functions. How this might be achieved at the cellular and molecular levels remains unknown, as ACh exerts broad and complex effects on various cell types within the DG (Pabst *et al*., 2016). To address this question, we took an unbiased approach to address cholinergic circuit activity-dependent changes in a cell-type-specific manner. Single-nucleus RNA-seq of the DG from mice with cholinergic circuit activation (ChR2) or sham stimulation controls (YFP) was performed to address cholinergic activity-induced transcriptional changes across various DG cell types. We adopted Split Pool Ligation-based Transcriptome sequencing (SPLiT-seq) to investigate transcriptomic changes in diverse DG cell types in response to cholinergic circuit activity (Figure 5A). Dorsal DGs from a total of 10 animals were micro-dissected and >20k cells passed quality control (5 animals for stimulated YFP control 10,072 cells; 5 animals for cholinergic circuit activation: 11,575 cells) (Figure S5). Following SPLiT-seq, we defined unbiased cell clusters via Louvain-Jaccard clustering with multi-level refinement and created a Uniform Manifold Approximation and Projection (UMAP) embedding to visualize the high-dimensional data in a two-dimensional projection (Figure S6B). To resolve the cell-type identities, we explored cluster-specific gene signatures and compared them to known signatures of hippocampal/DG cell types derived from single-cell/nucleus RNA-seq (Arneson et al., 2018; Artegiani et al., 2017; Ding et al., 2020; Habib et al., 2016; Hochgerner et al., 2018). We recovered all known major DG cell types, including mature granule cells (GCs) expressing *Prox1*, GABA interneurons (GABA) expressing *Gad1*/*Gad2*, mossy cells (MCs) expressing *Sv2b*, *Grm8*, or *Prrx17*, astrocytes expressing *Gfap* and *S100B*, oligodendrocytes (oligo) expressing *Mobp*, oligodendrocyte progenitor cells (OPCs) expressing *Pdgfra*, ependymal cells expressing *Foxj1*, microglia expressing *Cst3*, and endothelial cells expressing *Chd5* and *Flt1*. We also recovered cell clusters related to adult hippocampal neurogenesis, including rNSCs expressing *Hopx*, *Clu*, and *Aldoc*, (Artegiani *et al*., 2017; Shin et al., 2015), neural progenitors/neuroblasts (NPs/NBs) expressing *Ccnd2*, *Bhlhe22*, and *Gabra5* (Hochgerner *et al*., 2018) (Figure 5B, Figure S6, Figure S7, Figure S8B), and immature GCs expressing *Dsp* (Hochgerner *et al*., 2018). Our clusters exhibited consistent proportions among biological replicates, and cell classes corresponded to observed hierarchical clustering in our pairwise Pearson correlation coefficients plot (Figure 5C). Based on distinct marker gene expression within each major cell class, we identified that cell classes had distinct subclusters. The cell classes with multiple annotated subclusters included GCs (14 clusters), interneurons (5 clusters), and MCs (3 clusters) (Figure 5B and 5C). These results highlight the heterogeneity within these cell populations. In addition, we annotated 6 cornu ammonis (CA) clusters, 5 subiculum clusters, 1 ependymal cluster, and 1 Cajal-Retzius cluster. These cell clusters likely resulted from contamination (∼20% of total cells) during DG micro-dissection, so they were excluded from subsequent analyses.

**Figure 5:**
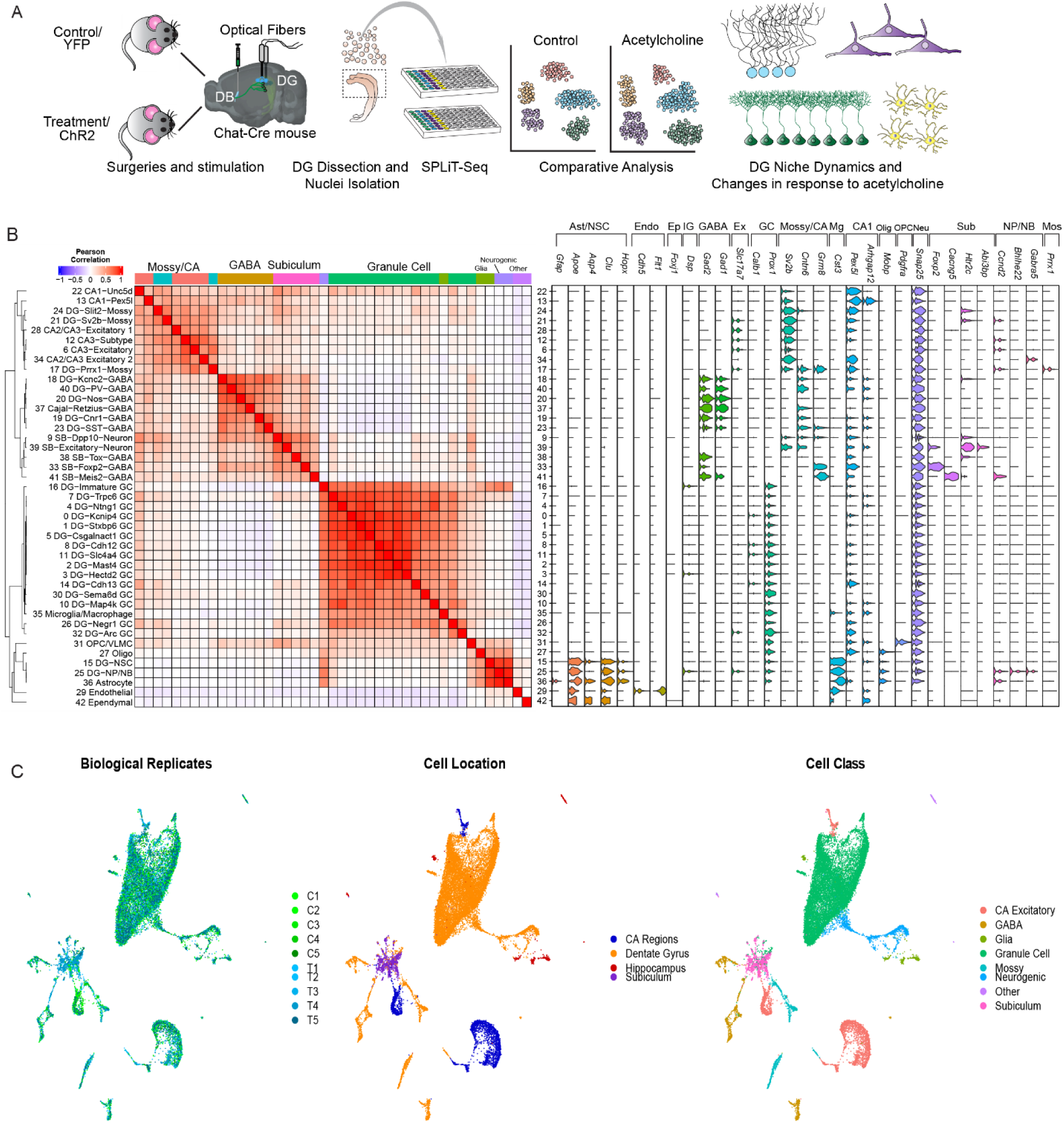
Identification of DG cells using snRNA-seq. A) Experimental schematic for snRNA-seq experiment upon circuit-specific stimulation of DB cholinergic afferents. B) Pearson pairwise correlation plot using the mean expression of the top fifty most representative genes for each cluster and a violin plot showing expression of marker genes within each cluster. Violin plot has labels for each cluster above marker genes that correspond to the following: Ast/NSC - Astrocyte/Neural Stem Cell, Endo – Endothelial, Ep – Ependymal, IG – Immature Granule Cell, GABA – GABA interneuron, Ex – Excitatory Neuron, GC – Granule cell, Mossy/CA – Mossy cells/ Cornu ammonis, Mg – Microglia, CA1 – Cornu ammonis 1, Olg – Oligodendrocyte, OPC - Oligodendrocyte Progenitor Cell, Neu – Neuron, Sub -Subiculum, NP/NB – Neural Progenitor/Neuroblast, Mos – Mossy Cells. C) Dimension reduction of clusters using UMAP plots to show clusters separated by biological replicates, cell location, or cell class.

### Cholinergic activity-dependent transcriptomic changes in DG mature neurons relate to synaptic functions critical for learning and memory

We next sought to address how activation of the DB-DG cholinergic circuit alters the transcriptional landscape of the DG across various cell clusters. We identified a total of 2,676 differentially expressed genes (DEGs) in response to cholinergic circuit activation across various types of DG cells (Wilcoxon rank-sum test p < 0.01, log_10_-fold change > 0.1), including diverse mature neuron populations, glia, endothelial cells, as well as adult-born cells within the neurogenic lineage (Figure 6A and 6B, Table S1). The majority of DEGs were distributed in mature neurons (2,006 out of 2,676 DEGs, 75%), including GCs (1,070 DEGs), interneurons (574 DEGs), and MCs (362 DEGs). Interestingly, cholinergic circuit activity differentially altered gene expression across distinct cell clusters within the same cell class (Figure 6A and 6B). For instance, the two GC clusters with the most DEGs, *Trpc6*-GC and *Kcnip4*-GC, only shared ∼10% common DEGs, suggesting that cholinergic activity differentially regulates distinct subtypes of GCs. Furthermore, many top-ranked DEGs in the neuronal clusters were involved in the regulation of synaptic transmission, calcium signaling, and synaptic formation (Figure 6C). For instance, the *Trpc6-*GC cluster showed significant upregulation of *Camk2d*, a calcium-dependent protein kinase required for memory formation (Zalcman et al., 2019). The *Kcnip4*-GC cluster exhibited significant upregulation of *Tenm2*, a tenurin protein that mediates calcium signaling using intracellular stores (Silva et al., 2011). The *Kcnc2*-GABA cluster (with the most DEGs within the interneuron population) exhibited significant upregulation of *Cacna1c*, a voltage-gated calcium channel that controls gene expression through coupling membrane depolarization with cAMP response element-binding protein phosphorylation (Wheeler et al., 2012). The SST-GABA cluster exhibited significant upregulation of Camk1, a calcium protein kinase involved in synaptic plasticity and neuronal survival (Bito and Takemoto-Kimura, 2003). The *Prrx1*-Mossy cluster (with the most DEGs within the MC population) exhibited significant upregulation of *Gpc4*, a heparan sulfate proteoglycan protein that regulates axon guidance and excitatory synapse formation (Ma et al., 2022). To validate some of the DEGs, we performed immunohistology of *Cdh13*, a cell adhesion protein involved in synaptic function necessary for learning and memory (Rivero et al., 2015). *Cdh13* was one of the most significantly upregulated genes in several GC clusters. Our immunohistology confirmed increased expression of *Cdh13* in GCs from circuit-stimulated mice as compared to the YFP controls (Figure 6D and 6E).

**Figure 6:**
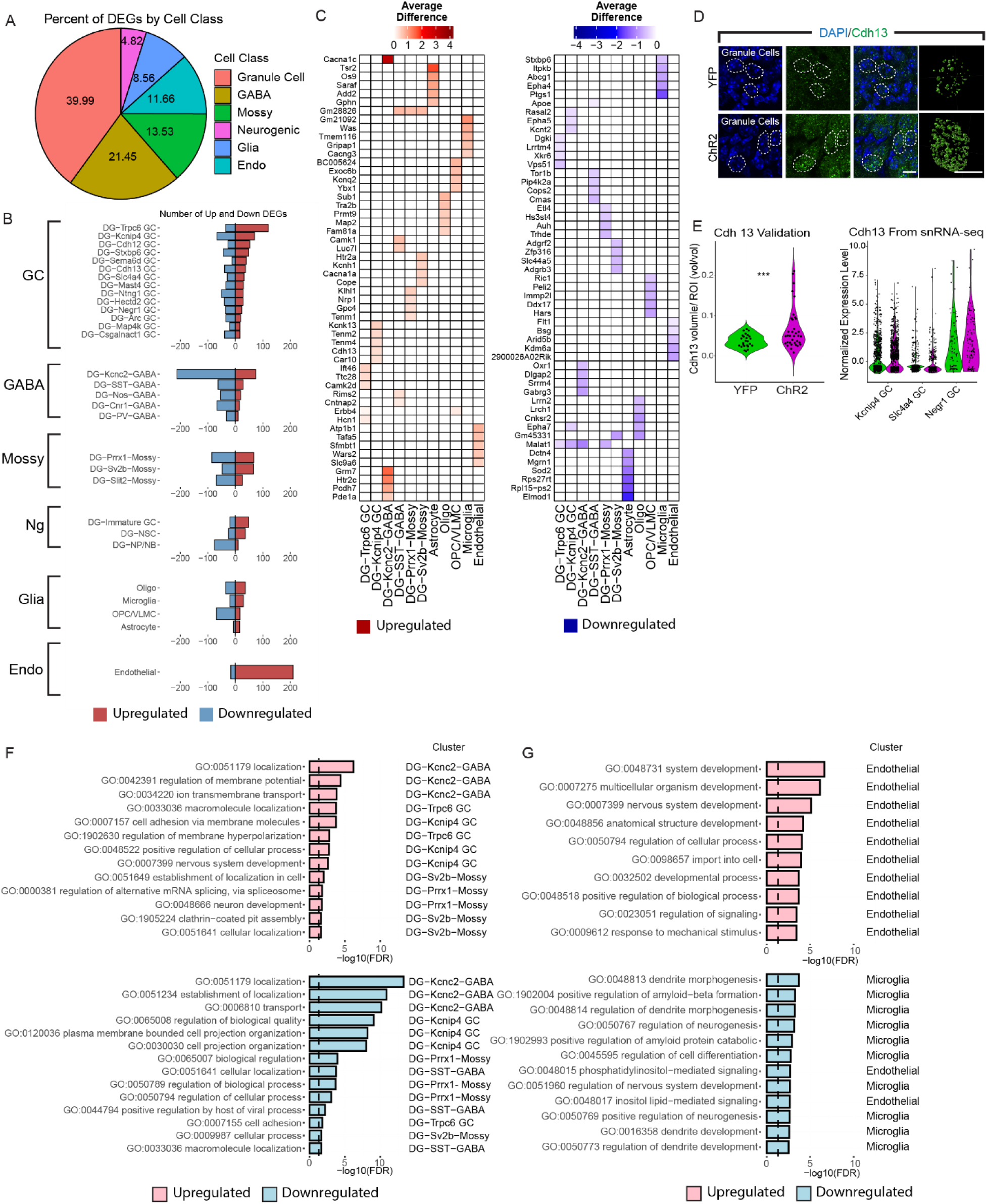
Cell-type specific transcriptomic changes in response to cholinergic circuit stimulation. A) Pie chart showing the percentage of DEGs by cell class. DEGs are counted more than once if they are repeated in multiple clusters within each cell class or within multiple clusters across different cell classes. B) Bar plots showing upregulated and downregulated DEGs in each cluster sorted by most upregulated DEGs. Analysis excludes subiculum and CA clusters since they aren’t part of the DG. Clusters are grouped by cell classes: GC – Granule Cell, GABA – GABA, Mossy – Mossy Cells, Ng – Neurogenic, Glia – Glia, Other – Cell types that don’t fall in previous categories. C) Heatmap showing top 5 most significant upregulated and downregulated DEG’s in the top 2 most responsive clusters for each cell class, except glia which includes astrocytes. D) Representative confocal image of Cdh13 immunofluorescence experiment with volumetric analysis. Confocal Scale bar = 20 µm Volumetric analysis scale bar = 2.5 µm E) Plot shows sum of all volumes within each cell. n = (25,35) cells from 3 biological replicates per condition. p-val = .0023 student’s t-test and clusters which have differentially expressed levels of Cdh13. F) Most significant GO terms from the top 2 most responsive neuronal clusters. Upregulated (pink) and down-regulated (light blue) with -log_10_(FDR) > 1.5. G) Most significant GO terms from most responsive glial cells that were upregulated (pink) and down-regulated (light blue) with -log_10_(FDR) > 1.5.

Furthermore, we performed gene ontology analyses of all the neuronal clusters (Figure S9). The neuronal clusters with most DEGs (top 2 clusters of each cell class) revealed upregulated GO terms involved in localization/macromolecule localization (such as protein localization to the cell surface and axons), regulation of membrane potential, ion transmembrane transport, and cell-cell adhesion via membrane adhesion molecules. Downregulated GO terms include localization/establishment of localization and transport (such as neurotransmitter uptake/secretion and synaptic vesicle transport), regulation of biological quality (such as regulation of neurotransmitter levels and calcium ion transport into the cytosol), and cell projection organization (such as neuron projection organization and pre/post synaptic membrane assembly) (Figure 6F). Together, these results revealed molecular mechanisms underlying cholinergic activity-dependent regulation of all the major neuronal types/subtypes in the DG. Importantly, these molecular changes collectively support cell-type/subtype specific synaptic remodeling in response to cholinergic circuit activity, which may be relevant for spatial memory encoding.

In contrast to neuronal cells, non-neuronal cells including glia and endothelial cells exhibited much fewer DEGs in response to cholinergic circuit activation (670 out of 2,676 DEGs, 25%) (Figure 6A and 6B). Gene ontology analyses revealed several significant biological pathways within endothelial cells and microglia. No significant GO terms were identified in astrocytes, oligodendrocytes, and oligodendrocyte progenitor cells. Interestingly, microglia exhibited downregulation of a GO term for positive regulation of amyloid-beta formation, which aligns with the established anti-inflammatory role of Ach (Patel et al., 2017; Rosas-Ballina and Tracey, 2009). Notably, DEGs in both endothelial cells and microglia exhibited GO terms primarily involved in developmental-like processes (Figure 6G, Table S2). Specifically, endothelial cells exhibited upregulated GO terms in nervous system and structural development, while microglia exhibited downregulated GO terms involved in dendritic morphogenesis and development, differentiation, and neurogenesis (Figure 6G). These results suggest that non-neuronal cells recruited by cholinergic circuit activation may play a critical role in regulating developmental processes with limited involvement in regulating synaptic functions.

### Cholinergic activity-dependent transcriptomic changes in rNSCs support neurogenic proliferation and structural development

Following the identification of neuronal DEGs, we examined cholinergic activity-induced DEGs in rNSCs. To confirm the lineage relationship of rNSCs with newborn progeny, we performed pseudotime analysis using Slingshot (Street et al., 2018). Our analysis revealed transitions from our rNSC cluster to the NP/NB cluster and then to the immature GC cluster and finally to the *Mast4*+ GC cluster (Figure 7A and 7B). These results recapitulated the known lineage progression of these clusters during adult hippocampal neurogenesis. Interestingly, when comparing genes in rNSCs versus NPs/NBs, we identified increased expression of *Mertk* in NSCs, a TAM receptor that supports neural stem cell survival and proliferation (Ji et al., 2014), and increased expression of *Calm1* and *Calm2*, two calmodulin proteins that aid neuroblast migration in the NP/NB cluster (Khodosevich et al., 2009) (Figure 7C). Together, these results further confirmed the classification of our clusters along a developmental lineage and supported our identification of the rNSC cluster.

**Figure 7:**
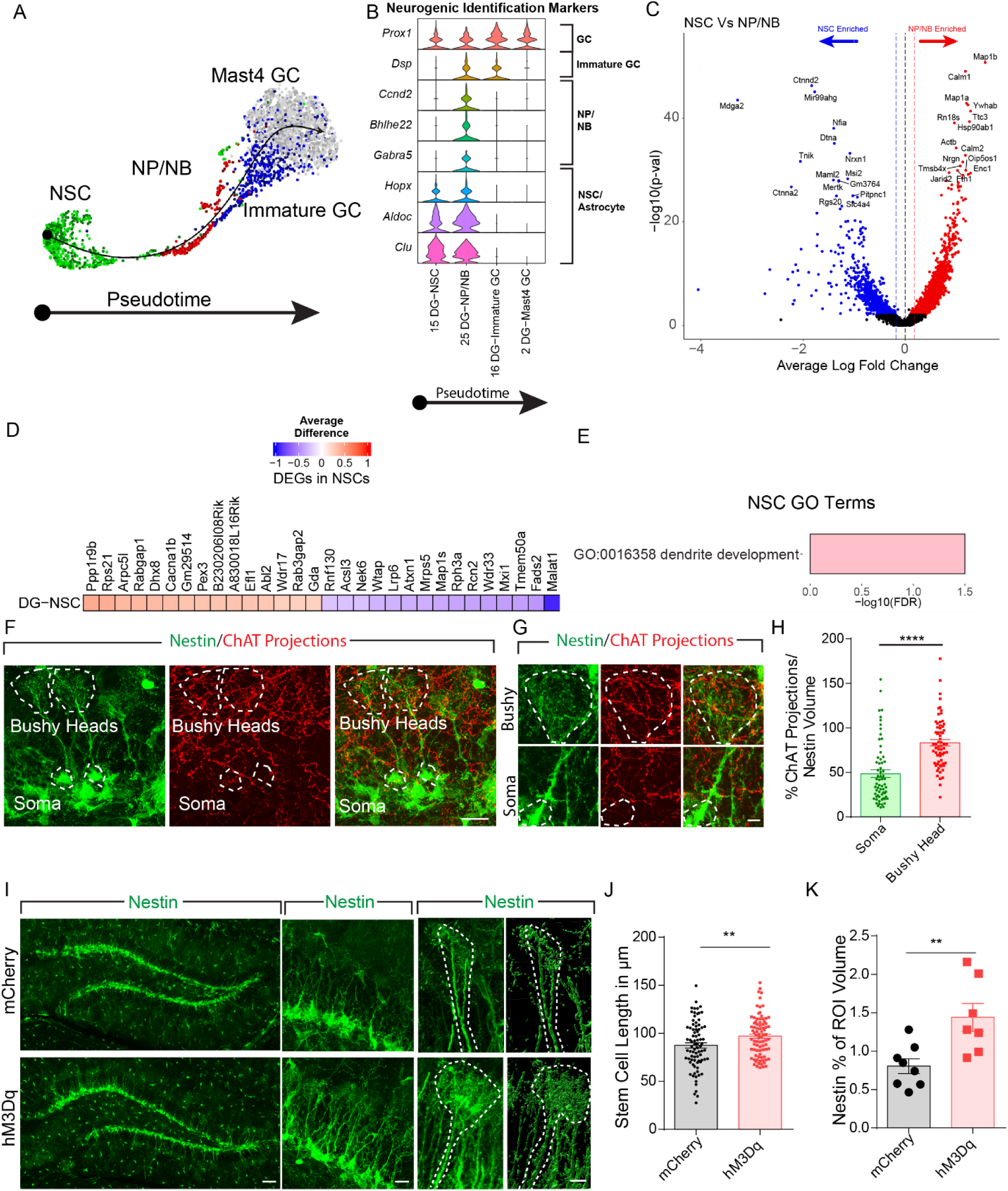
Cholinergic activity-dependent transcriptomic changes in rNSCs support their neurogenic proliferation and structural development. A) Slingshot pseudotime analysis of clusters from the neurogenic cell class and the most proximal GC cluster. B) Violin plots showing marker gene expression in neurogenic cell class clusters. Markers are taken from (Artegiani *et al*., 2017; Hochgerner *et al*., 2018; Shin *et al*., 2015). C) Volcano plot showing differentially upregulated genes in the combined, both YFP (control) and ChR2 (treatment) rNSC cluster (red) and NP/NB cluster (blue). D) Top 10 most significant upregulated and downregulated DEGs in the rNSC cluster. E) Go term associated with DEGs in our rNSC cluster. F) Representative image of rNSC bushy heads and Soma interacting with cholinergic fibers from the DB. Scale bar = 20 µm G) Close-up image of rNSC bushy head and soma interacting with cholinergic fibers. Scale bar = 5 µm. H) Quantification of cholinergic projections with Soma and Bushy Heads. Bars indicate mean +/- S.E.M. n = 65, p-val <.0001 by paired student’s t-test. I) Representative image of rNSCs from a ChAT-Cre::Nestin:GFP animal who was chemogenetically stimulated in DB nuclei. Left scale bar = 20 µm, middle scale bar = 5 µm, right scale bar = 2 µm. J) Quantification of stem cell length in mCherry and chemogenetically stimulated groups. Bars indicate mean +/- S.E.M. n = (82,85), p-val = .0073 student’s t-test. K) Quantification of bushy heads as a percent of ROIs within DG in control or hM3Dq animals. Bars indicate mean +/- S.E.M., n = (8,6), p-val = .007 student’s t-test.

Cholinergic circuit activation induced a total of 61 DEGs (36 up, 25 down) in the rNSC cluster (Figure 6B and Figure 7D). Specifically, the rNSC cluster exhibited DEGs associated with the generation of new neurons (*Atxn1, Mapk8, Atp7a, Acsl3, Tenm2*) and actin-mediated structural development (*Abi1, Abl2, Mapk8, Atp7a*). Our functional studies have shown that both optogenetic activation and chemo-activation of DB cholinergic neurons resulted in a significant increase of proliferating rNSCs (Figure 2 and Figure S2), thus supporting the neurogenic function of rNSCs. In addition, gene ontology analyses revealed GO terms involved in the development of processes of rNSCs, suggesting that actin-related function in rNSCs may be involved in regulating the structure of rNSCs. The rNSCs possess prominent radial processes that terminate as a bushy head. Importantly, these processes were found to be associated with DB-DG cholinergic projections (Figure 7F). Interestingly, radial processes and bushy heads of rNSCs exhibited more associations with DB-DG cholinergic projections than the soma of rNSCs (Figure 7H), suggesting that they may serve as a critical functional domain in response to cholinergic circuit activity. To test this, we examined the length of the radial processes and the size of the bushy heads of rNSCs after chemogenetic activation of DB cholinergic neurons using ChAT-Cre::Nestin:GFP mice injected with AAV-DIO-hM3Dq-mCherry or AAV-DIO-mCherry in the DB (Figure 7I). Strikingly, activation of DB cholinergic neurons led to both increased length of the radial processes and increased size of bushy heads in rNSCs (Figure 7J and Figure 7K), thus supporting cholinergic activity-dependent structural remodeling of the rNSCs. Together, these results provided the molecular mechanisms to support the possibility that radial/bushy processes of rNSCs couple the cholinergic circuit activity to the neurogenic proliferation of rNSCs.

### Cholinergic circuits recruit mature granule cells to impact rNSC activity

Our snRNA-seq data revealed significant upregulation of genes involved in regulating the activity of rNSCs, such as *Cacna1b* (voltage-gated calcium channel subunit Alpha1 B) (Figure 7D). Therefore, we wondered whether activation of the DB-DG cholinergic circuit impacts the activity of rNSCs, and if so, whether this is through direct or indirect mechanisms. To address this question, we performed slice electrophysiology in ChAT-Cre::Nestin:GFP mice injected with AAV-DIO-ChR2 in the DB. GFP+ rNSCs were recorded for light-evoked currents upon optogenetic activation of DB-DG cholinergic projections (8Hz, 5ms, 2s) (Figure 8A). Interestingly, optogenetic activation of DB-DG projections induced inward currents in a quarter of recorded rNSCs (5/21) (Figure 8B and 8C). It has been shown that rNSCs respond to bath application of ACh with increased calcium activity through M1 receptors (Itou et al., 2011). However, we found these light-evoked inward currents were not affected by atropine, a muscarinic ACh receptor antagonist (Figure 8D), suggesting an indirect mechanism. Unexpectedly, these light-evoked inward currents were largely blocked by ionotropic glutamate receptor (iGluR) antagonists AP5 (for NMDARs) and CNQX (for AMPARs) (Figure 8E), suggesting that iGluRs (but not mAChRs) mediate cholinergic activity-induced depolarization of rNSCs. These results aligned with our recent studies showing rNSC depolarization upon activation of glutamate-releasing niche cells through iGluR-mediated signaling (Asrican *et al*., 2020; Yeh *et al*., 2018). These results indicated that the DB-DG cholinergic circuit requires a glutamatergic intermediary to relay signals to rNSCs.

**Figure 8:**
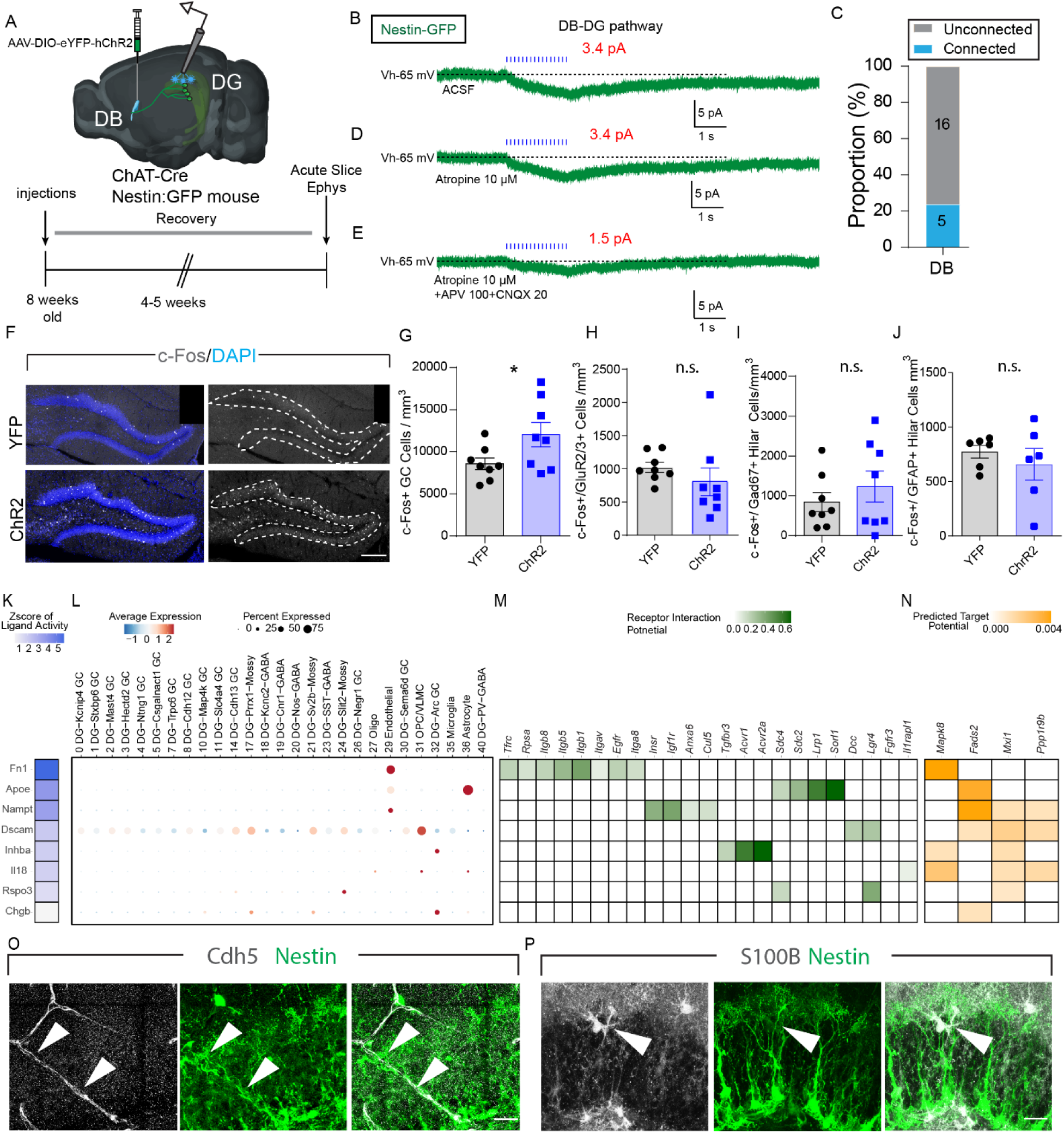
Cholinergic circuits regulate rNSCs through granule cells, endothelial cells, and astrocytes. A) Schematic of acute hippocampal slice electrophysiology experiment to record from rNSCs after stimulation of cholinergic afferents using optogenetics. B) Voltage clamp recordings of Nestin:GFP+ rNSCs in the presence of ACSF, (muscarinic antagonist) atropine 10 µm, or atropine 10 µm and (NMDA antagonist) APV and (AMPA antagonist) CNQx 20 µm after optogenetic stimulation. C) Quantification of connected rNSCs after optogenetic stimulation from acute slice experiments. F) Representative image of DG sections quantified for C-Fos expression after DB optogenetic stimulation. Scale bar = 100 µm. G) Quantification of c-Fos in Granule cells, bars indicate mean +/- S.E.M. n= (8,8), p-val =.0348 student’s t-test. H) Mossy cells, I) Gaba Cells J) Astrocytes n= (6,6). K) Nichenet predicted ligands ordered by Z-Score taken from Pearson’s Correlation. L) Dot plot showing expression of ligands in sender clusters used to generate predictions about receptor and target gene interactions. M) Predicted ligand receptors within rNSCs from snRNA expression data using Nichenet. N) Predicted target genes within rNSCs from snRNA expression data using Nichenet. O) Image showing endothelial cells interacting with rNSCs within the DG. Scalebar = 20 µm P) Image of astrocytes interacting with rNSCs in the DG. Scalebar = 20 µm

We next sought to address the niche cell type(s) activated by stimulating cholinergic circuits to mediate cholinergic activity-dependent regulation of rNSCs. We examined c-Fos expression in GCs, MCs, interneurons, and astrocytes, after cholinergic circuit activation. Interestingly, a significant increase was observed in the density of c-Fos+ GCs (Figure 8F and 8G). This finding is consistent with recent reports that stimulating cholinergic afferents lowed the threshold for GC firing thus making them more excitable, potentially through interneuron-mediated disinhibition (Martinello et al., 2015; Ogando et al., 2021). By contrast, no significant difference was observed in the density of c-Fos+ GluR2/3+ MCs (Fig. 8H), c-Fos+ GAD67+ interneurons (Figure 8I), and c-Fos+ GFAP+ astrocytes (Figure 8J). These results suggest that cholinergic activity-dependent glutamate transmission from GCs is likely to be the glutamatergic source for rNSCs. Supporting this notion, electron microscopy showed radial processes of rNSCs wrapped around dendritic spines of mature GCs (Moss et al., 2016). Along with cholinergic activity-induced structural changes in radial/bushy processes of rNSCs (Figure 7J and 7K), these findings suggested that radial processes/bushy heads of rNSCs could sense cholinergic activity-dependent glutamate signaling from dendritic spines of the GCs.

### NicheNet analyses reveal endothelial cells and astrocytes as additional intermediates for cholinergic circuit activity-dependent regulation of rNSCs

The unexpected finding that cholinergic circuit activity-induced depolarization of rNSCs requires a niche intermediary suggests that DG niche cells maybe more sensitive to cholinergic activity than rNSCs. Supporting this notion, our snRNA-seq results showed that multiple types of DG niche cells respond to cholinergic circuit activation with significantly more DEGs than rNSCs. These results promoted us to search for additional niche cells that may mediate cholinergic activity-dependent regulation of rNSCs but cannot be otherwise detected by electrophysiology.

In addition to the release of transmitters by neurons and astrocytes, various cell types in the brain can alter the expression of surface ligands or secrete chemical ligands in response to certain stimuli. Therefore, cholinergic circuit activation may alter the level of ligand expression/release from various DG niche cells to communicate with rNSCs through distinct intracellular mechanisms. To predict potential gene targets of intercellular communication between niche cells and rNSCs, we adopted NicheNet, a computational method that predicts ligand-target links between interacting cells by combining their expression data with existing knowledge on signaling and gene regulatory networks (Browaeys et al., 2020). The DEGs from our snRNA-seq data in both neuronal and non-neuronal clusters were included as niche cells for rNSCs in an unbiased manner. Our NicheNet analysis identified a total of 8 prioritized ligands, 25 potential receptors, and 4 intracellular downstream genes from our niche cells and rNSCs. Notably, Fibronectin 1 (FN1) from endothelial cells and Apolipoprotein E (ApoE) from astrocytes were identified as the top-ranked ligands (Figure 8K and 8L). FN1 is an extracellular matrix (ECM) glycoprotein that binds integrins Itgb5 and Itgb1 in rNSCs (Figure 8M) and regulates Mapk8 intracellularly (Figure 8N); and ApoE from astrocytes binds Sorl1 and Lrp1 in rNSCs and regulates Fads2 intracellularly (Figure 8K - 8N). Recent studies showed that FN1 and ApoE serve as critical niche signaling molecules to regulate adult rNSC quiescence and maintenance (Morante-Redolat and Porlan, 2019; Tensaouti et al., 2018; Yang et al., 2011). These findings support their roles in regulating adult rNSCs.

Supporting the functional interaction between endothelial cells/astrocytes and rNSCs, we found close morphological associations between these niche cells and rNSCs (Figure 8L and 8M). Taken together, these results suggest a complementary mechanism by which cholinergic circuits may recruit non-neuronal niche cells such as endothelial cells and astrocytes to interact with rNSCs through surface/secretory ligands such as FN1 or ApoE. This mechanism mediated by non-neuronal cells may coexist with the GC-mediated mechanism to collectively regulate rNSCs in response to cholinergic circuit activity.

## Discussion

Although much is known about the role of Ach as a modulator of information flow in hippocampal CA1 and CA3 subregions (Hasselmo, 2006; Hasselmo and Schnell, 1994; Palacios-Filardo et al., 2021; Vogt and Regehr, 2001), cholinergic modulation of the DG microcircuit in the contexts of learning/memory and adult neurogenesis has been less explored. Furthermore, the major cholinergic projections from basal forebrain to the hippocampus have been studied by targeting combined medial septum (MS) and DB, therefore, the relative contribution of DB-DG versus MS-DG cholinergic inputs in regulating DG functions remains elusive. To fill these knowledge gaps, our study demonstrated that activation of DB-DG cholinergic projections promotes quiescent rNSC activation and spatial memory. In addition, most studies on cholinergic modulation of the DG were performed using slice electrophysiology (Martinello *et al*., 2015; Ogando *et al*., 2021; Pabst *et al*., 2016). Although these studies have undoubtedly provided valuable information on cholinergic regulation of specific types of DG cells, cell-type-specific signaling mechanisms underlying cholinergic activity-dependent modulation of DG is largely lacking. To address how cholinergic circuits modulate diverse yet distinct DG cells in an unbiased fashion, we performed single-nucleus RNA-seq of DG and identified broad transcriptomic changes across multiple types of mature and adult-born cells in response to cholinergic circuit activation. Notably, we found cholinergic-activity-induced molecular changes related to synaptic functions crucial for spatial memory in various neuronal populations, including GCs, interneurons, and MCs, as well as changes related to structural development and proliferation of rNSCs. It is worth noting that while single-cell/nucleus RNA-sequencing has been extensively performed in various brain regions including the DG, our study made the first attempts to profile transcriptomic changes induced by specific neural circuits at the single-nucleus level in a highly heterogenous brain structure. Given that cholinergic modulation exerts broad actions on multiple types of brain cells and not all these actions are beneficial to brain functions, the knowledge gained from our studies will guide future therapeutic strategies by selectively targeting cell-(sub)type-specific molecular components that are beneficial to circuit and brain functions.

One unexpected finding is that cholinergic circuit activity-induced depolarization of rNSCs requires a glutamatergic intermediary, likely GCs. This is in sharp contrast to adult-born immature neurons that have been shown to receive direct DB cholinergic inputs through M1 muscarinic ACh receptors (Zhu et al., 2017). The failure to receive direct cholinergic signals in rNSCs could be explained by the low expression level of ACh receptors in rNSCs (Shin *et al*., 2015). This study adds another example to our current evidence supporting the critical role of glutamate-releasing niche cells in regulating rNSCs. Specifically, glutamate-releasing niche cells either directly signal rNSCs, such as mossy cells (Yeh *et al*., 2018), or serve as a relay to mediate signals from other cells, such as astrocytes (Asrican *et al*., 2020; Pabst *et al*., 2016). These studies support the general principle that rNSCs are highly responsive to glutamatergic signaling in the DG neurogenic niche. This raises the question of how rNSCs sense glutamate signaling. Our snRNA-seq data revealed cholinergic activity-induced DEGs associated with the development of radial processes/bushy heads of rNSCs. Our recent electron microscopy analyses showed that bushy heads of the rNSCs express NMDA receptors and wrap around glutamatergic synapses (Yeh *et al*., 2018). These results suggest that the bushy heads of rNSCs might be the glutamate-sensing site, which can potentially signal the soma of the rNSCs to induce membrane depolarization and neurogenic proliferation of rNSCs. Future studies expressing genetically-encoded calcium or glutamate sensor in rNSCs in combination with cholinergic circuit stimulation will be able to address this question.

In addition to glutamate-releasing cells (likely GCs), we performed NicheNet analyses to identify additional relay niche cells that could not be otherwise detected by slice electrophysiology. We are the first to adopt NicheNet analyses in the context of circuit-activity-dependent niche regulation of adult rNSCs. Using this platform, we identified novel niche cells and their ligands that interact with rNSCs though receptor-mediated intracellular signaling. FN1 from endothelial cells and ApoE from astrocytes were identified as top candidates that can potentially serve as intermediaries for cholinergic activity-dependent regulation of rNSCs. It has been shown that the adult subependymal zone (SEZ) is rich in ECM molecules, such as fibronectin, and brain endothelial cell-derived ECM via integrins in the SEZ regulates quiescence and proliferation of adult NSCs (Hirano and Suzuki, 2019; Ottone et al., 2014; Porcheri et al., 2014). Despite the established role of ECM molecules in regulating adult SEZ NSCs and the close anatomical associations between vasculature and rNSCs in the DG, the activity-dependent role of endothelial cell-derived ECM-integrin signaling in rNSC regulation remains to be established. A recent study showed that neural circuit activity from interneurons can affect the blood flow, which in turn impacts adult hippocampal neurogenesis (Shen *et al*., 2019). This study supports activity-dependent neurovascular coupling in the regulation of adult neurogenesis. In contrast to the role of FN1, ApoE is critical for rNSC maintenance (Yang *et al*., 2011). Therefore, FN1 and ApoE may act in concert to regulate the balance of rNSC proliferation and pool maintenance in response to cholinergic circuit activation. Taken together, our findings suggest that the DB-DG cholinergic circuit appears to preferentially recruit DG neuronal populations to regulate synaptic functions that support spatial memory, while preferentially recruiting DG non-neuronal populations to regulate rNSCs and neurogenesis (Figure S10). Interestingly, cholinergic circuit-recruited GCs appear to play dual roles to regulate both spatial memory and rNSCs. These findings fill the gap in our understanding of how neural circuits orchestrate complex signaling from diverse DG populations to support distinct hippocampal functions. These studies also spark our future efforts to validate these novel candidates in regulating rNSCs and hippocampal neurogenesis.

Deficits in cholinergic signaling are highly implicated in Alzheimer’s disease (AD) (Kihara and Shimohama, 2004; Muir, 1997; Perry, 1988). Both spatial memory loss and reduced adult hippocampal neurogenesis have been reported in AD patients (Moreno-Jimenez et al., 2019; Selkoe, 2002). Our studies showed that activation of the DB-DG cholinergic circuit promotes both spatial memory and rNSC activation, thus posing the possibility that stimulating cholinergic circuits may restore spatial memory and hippocampal neurogenesis in AD patients. In addition, deposition of Aβ plaques is a pathological hallmark in AD brains. Interestingly, our snRNA-seq data suggested that microglia exhibit downregulation of biological processes related to Aβ formation in response to cholinergic circuit activation, suggesting that cholinergic activity potentially reduces plaque deposition in AD through a microglia-mediated mechanism to prevent plaque formation. Therefore, exploring the therapeutic potential of the DB-DG cholinergic circuit to treat neurodegenerative diseases such as AD may represent a novel therapeutic strategy.

## Acknowledgments

We thank all the members of the Song lab for comments and discussions. This work was supported by grants from NIH (MH111773, MH122692, AG058160, NS104530 to J. Song, R35NS116843 to H. Song, and R35NS097370 to G-l.M.) and Alzheimer’s Association to J. Song. L. Quintanilla was supported by a NIA Ruth L Ruth L Kirschstein F31 predoctoral fellowship (AG067718) and a NIMH R01 Diversity Supplement (3R01MH111773-03S1). Confocal microscopy was performed at the UNC Neuroscience Microscopy Core Facility (RRID: SCR_019060) with the technical assistance from Dr. Michelle S Itano. The Neuroscience Microscopy Core was supported in part by funding from the NIH-NINDS Neuroscience Center Support Grant P30 NS045892 and the NIH-NICHD Intellectual and Developmental Disabilities Research Center Support Grant U54 HD079124.

## Author contributions

J. Song supervised the project, designed the experiments, and wrote the paper. L. Quintanilla designed the experiments, wrote the paper, and carried out all the experiments and data analysis. Y. Su, G-l. Ming, and H. Song performed single-cell sequencing experiment. J. Simon helped RNA-Seq analysis. Y-J Luo and B. Asrican carried out slice electrophysiology experiments. S. Tart, R. N. Sheehy, and Y-D Li assisted some aspects of this study.

## Declaration of Interests

G-l. Ming is on the advisory broad of Cell Stem Cell. The authors declare no competing interests.

## KEY RESOURCES TABLE

| REAGENT or RESOURCE | SOURCE | IDENTIFIER |
| --- | --- | --- |
| **Antibodies** |  |  |
| rabbit anti-RFP | Rockland | Cat# M11217, RRID: AB_2536611 |
| goat anti-ChAT | Millipore | Cat# AB144P, RRID: AB_2079751 |
| rabbit anti-cFos | SySy | Cat# 226003, RRID: AB_2231974 |
| chicken anti-nestin | Aves | Cat# NES, RRID: AB_2314882 |
| mouse anti-Chd5 | RND Systems | Cat# AF1002-SP, RRID: AB_2077789 |
| mouse anti-GFAP | Millipore | Cat# MAB360, RRID: AB_11212597 |
| mouse anti-Gad67 | Millipore | Cat# MAB5406, RRID: AB_2278725 |
| goat anti-GFP | Rockland | Cat# 600-101-215, AB_218182 |
| rabbit anti GluR2/3 | Sigma | Cat# AB1506, RRID: AB_90710 |
| mouse anti-S100B | Atlas Antibodies | Cat#AMAb91038, RRID: AB_2665776 |
| **Bacterial and Virus Strains** |  |  |
| AAV5-ef1a-DIO-hChR2-eYFP | UNC Vector Core | NA |
| AAV5-ef1a-DIO-eYFP | UNC Vector Core | NA |
| AAV5-hsyn-DIO-hM3Dq-mCherry | Addgene | RRID: Addgene_44361 |
| AAV5-hsyn-DIO-mCherry | Addgene | RRID: Addgene_50459 |
| AAV5-flex-taCasp3-TEVp | UNC Vector Core | NA |
| **Chemicals, Peptides, and Recombinant Proteins** |  |  |
| bicuculline | Tocris | 0131 |
| NBQX | Tocris | 0373 |
| APV | Tocris | 0105 |
| TTX | Tocris | 1078 |
| AIDA | Tocris | 0904 |
| AMPA | Tocris | 0254 |
| NMDA | Tocris | 0114 |
| DHPG | Tocris | 0805 |
| Carbachol | Tocris | 2810 |
| EdU | Carbosynth | NE08701 |
| CNO | NIH | C-929 |
| ω-agotoxin TK | Alamone Labs | STA-530 |
| **Critical Commercial Assays** |  |  |
| RNeasy mini kit | QIAGEN | cat. # 74104 |
| RNase-Free DNase Set | QIAGEN | cat. # 79254 |
| Click-iT_EdU Alexa Fluor_488 Imaging kit | ThermoFisher | cat. # C10337 |
| NEBNext_Ultra RNA Library Prep Kit for Illumina | New England Biolabs | cat. # E7530L |
| Nextera XT DNA Library Preparation Kit | Illumina | cat. # FC-131-1024 |
| Dynabeads MyOne Streptavidin C1 Kit | Thermo Fisher cat. | cat. # 65001 |
| Kapa HiFi HotStart Master Mix | KAPA Biosystems | cat. # 65001<br>Kapa HiFi HotStart Master Mix<br>KAPA Biosystems cat. # KK2600 |
| Deposited Data |  |  |
| Raw and analyzed snRNA-seq Data | This paper | GEO: GSE202481 |
| Analysis Codes | This Paper | Github:<br>https://github.com/lquin003/Song-Lab-Quintanilla-et-al |
| Experimental Models: Organisms/Strains |  |  |
| Mouse: C57BL/6 | Jackson Laboratory | RRID: IMSR_JAX:000664 |
| Mouse: ChAT-IRES-CRE | Jackson Laboratory | RRID: IMSR_JAX:006410 |
| Mouse: Nestin-GFP | Dr. Grigori Enikolopov at Stony Brook University | RRID: IMSR_JAX:033927 |
| Software and Algorithms |  |  |
| Imaris | Bitplane | RRID: SCR_007370 |
| FIJI | ImageJ | RRID: SCR_002285 |
| pClamp | Molecular Devices | RRID: SCR_011323 |
| Micro-Manager | Micro-manager.org | RRID: SCR_000415 |
| Igor Pro | Wavemetrics | RRID: SCR_000325 |
| Neuromatic | Thinkrandom.com | RRID: SCR_004186 |
| Olympus FluoView | Olympus | RRID: SCR_014215 |
| Zeiss ZEN | Zeiss | RRID: SCR_013672 |
| Prism | Graphpad | RRID: SCR_002798 |
| Adobe Illustrator | Adobe | www.adobe.com |
| Seurat V3 and V4 | (Stuart et al., 2019) | www.stijalab.org/seurat |
| zUMIs | Github | https://github.com/sdparekh/zUMIs |
| ggplot2 | Wickham 2016 | https://ggplot2.tidyverse.org |
| Slingshot | (Street et al. 2018) | https://github.com/kstreet13/slingshot |
| NicheNet | (Browaeys et al. 2019) | https://github.com/saeyslab/nichenetr |
| Complex Heatmap | Gu et al. 2016 | https://github.com/jokergoo/ComplexHeatmap |
| R, Rstudio V 4.1 | Rstudio | https://www.ncbi.nlm.nih.gov/pmc/articles/PMC4023662/ |

## Methods

### Animals

Single or double transgenic young adult mice (6-12 weeks, males and females) were used and randomly assigned to experimental groups. C57BL/6J, ChAT-IRES-Cre were obtained from Jackson laboratory stock #(006410), and Nestin:GFP mice (Mignone et al., 2004) were obtained from Dr. Grigori Enikolopov at Stony Brook University. All ChAT-IRES-Cre animals used in this study were heterozygous and came from breeding a homozygous ChAT-IRES-Cre animal with a wild type C57Bl/6J. Our double transgenic animals were generated by breeding a heterozygous Nestin:GFP animal with a homozygous ChAT-IRES-Cre animal. Animals were group-housed and bred in a dedicated husbandry facility with 12 hour /12 hour light-dark cycles with food and water ad libitum and received veterinary supervision. Animals that were exposed to surgical procedures were moved to a separate satellite housing facility for recovery with the same light-dark cycle. Animals were sacrificed at the same time for each experiment which was around 5:00 PM to 7:00 PM. No immune deficiencies or other health problems were observed in these mouse lines, and all animals were experimentally and drug-naive before use. All animal procedures followed the NIH Guide for Care and Use of Laboratory Animals and were approved by the Institutional Animal Care and Use Committee at the University of North Carolina at Chapel Hill.

### Stereotaxic Injections

Stereotaxic injections were performed on animals 6-8 weeks of age like that described in (Quintanilla et al., 2019). Briefly, animals were anesthetized using a 5% isoflurane/oxygen mix until animals were non-responsive to a toe pinch. Animals were then placed in a sterotax, eye ointment was applied, and the fur was removed using a hair removal product (Nair). The area was cleaned and local anesthetic, lidocaine ointment, was applied. The area was further cleaned using aseptic techniques and a 1-2 mm incision was created to expose the skull. The skull was cleaned to remove the connective tissue over the skull and small openings were created above injection sites using a 1mm microdrill. The following coordinates were used for injections: Dentate Gyrus A/P -2.00 mm, M/L +/- 1.5 mm, D/V -2.15 mm, Medial Septum A/P .75 mm, M/L 0.0 mm, D/V - 3.50, Diagonal Band A/P .75 mm, M/L +/-.60 mm, D/V -5.40 mm. A total of 400 nL of AAV virus was injected into either the DG, MS, or DB, with a flow rate of 75 nl/min for the DG or 100 nl/min for the MS/DB. After injection, the needle was left in the brain for a total of 5 minutes, to allow for the dispersal of AAV viral particles in the targeted brain region.

For optogenetic experiments we implanted optical fibers in the DG at the following coordinates: A/P -2.00 mm, M/L +/- 1.5mm, D/V -1.75 mm. Optical fibers were implanted after viral injections, and a head cap was created using dental cement. Dental cement was left to dry for at least 10 minutes until the animal was removed from the sterotax. After surgery, the animal was allowed to recover on a heated animal pad until normal locomotion resumed and awake behavior was visible. Postoperative care was provided by administering Meloxicam subcutaneously at pre-operation and 24-hour intervals. Animals were monitored by both veterinary staff and researchers during recovery to assure proper recovery until used for experiments.

### Tissue Processing

Animals were transcardially perfused with 4% PFA immediately after stimulation for neurogenesis and snRNA-seq experiments. Animals that were used for behavior had a wait time of 1 week after behavior before stimulation and perfusion for immunohistochemistry experiments to mitigate any residual behavior-related activity in the brain. After PFA fixation, brain tissue was cryopreserved in 30% sucrose for at least 2 days until ready for processing. The tissue was sliced using a microtome at 40 microns to collect both basal forebrain and hippocampus brain regions. A total of 40 microns X 64 sections were collected spanning about 2.5 mm of the hippocampus. For the MS/DB, 40 microns X 48 sections were collected spanning a total of about 2.0mm. Tissue was stored in cryoprotectant antifreeze at -20 degrees until used for immunofluorescence experiments.

### Immunofluorescence Experiments

Immunofluorescence experiments were performed similar to (Quintanilla et al. 2019). For immunofluorescence experiments not requiring Nestin, the following steps were taken. Briefly, sliced brain sections were taken from cryoprotectant antifreeze solutions when ready and placed in TBS-Triton (.05% Triton in TBS) and washed twice for 5 minutes on a shaking rocker. Sections were then permeabilized using permeabilization buffer (.5% TBS-Triton) for 20-30 minutes on a shaker. Sections were then incubated in blocking buffer on a shaker (TBS + (5 % Donkey Serum) for 30 minutes. After blocking, primary antibodies were mixed with blocking buffer solution and applied to floating sections in a 24-well plate. Sections were incubated for 48 hours at 4C. The following primary antibodies were used at the following concentrations goat anti-GFP 1:1000 (Rockland), rabbit anti-RFP 1:1000 (Rockland), goat anti-ChAT 1:250, chicken anti-Nestin 1:200, rabbit anti-c-Fos 1:100, rabbit anti-Cdh-13 1:1000.

For neurogenesis experiments, we performed antigen retrieval to better visualize Nestin expression. Antigen retrieval consisted of mounting brain sections on a slide and allowing them to dry. They were then placed in a solution of 0.1M citrate buffer and kept at a temperature between 92-95C for 7 minutes using a microwave as described in (Quintanilla *et al*., 2019). Brain sections then remained in citrate buffer for approximately 1 hour and were allowed to cool down to room temperature. Brain sections were then permeabilized and blocked as described previously.

### Image acquisition using Confocal Microscopy

Images were obtained using an Olympus FV3000 microscope. For anterograde tracing experiments the entire DG’s were imaged using a 20x objective with 2x zoom, 1-micron step size, and 8x averaging using a resonant scanner. When quantifying adult neural stem cells, entire DGs were scanned with a Galvano scanner instead using a 20x objective with 2x zoom and 4x averaging and a 1 micron step size. For morphology analysis, images were taken with a 60x objective, 8x averaging, and 2x zoom with a 0.5-micron step. Representative images of Neural Stem cells were also taken with morphology analysis settings using a 60x objective with 8x averaging and 2x zoom.

### Image analysis

Imaging analysis was an essential component to understand several aspects of this study. Analysis was performed using two separate analysis software. The first analysis software used was the Fiji/ImageJ version that ran on Java 6. The second utilized Imaris, which ran on version 9.1.0. Imaris versions were updated regularly as new updates became available.

For anterograde tracing, we utilized Imaris to perform a volumetric analysis of the projections from either the MS or DB. A total of 5 sections/animal were used for a total of 3 animals. ROIs were manually drawn using hippocampal landmarks to distinguish between different regions. ROIs are shown in (Figure 1B and Figure S1B). Each hippocampal section had been immunofluorescence stained to enhance visualization of projections with fluorophore-specific antibodies. ROIs were the same for both MS and DB with the only difference being the projection volume within ROIs for each animal. The reported values are a measurement of the sum of the volume of all objects within each ROI divided by ROI volume to normalize for different size ROIs. Hippocampal sections spanned from the dorsal DG to the start of the Ventral DG.

For quantification of projections to rNSCs, we took higher resolution images at 60x oil lens with 2X zoom. Images were obtained using the Galvano scanner on the FV3000 with 4X averaging. RNSCs were then selected for individual analysis by drawing ROIs around the same stem cell’s soma and bushy head.

For neurogenesis experiments, images were manually quantified using FIJI by a second carefully trained researcher who was blind to all experimental conditions. This included quantifications of all stem cell-related figures (Figure 2, Figure 3, Figure S2, Figure S3). A total of 5 sections were quantified per animal spanning the same Dorsal to Ventral axis used for anterograde tracing analysis. Manual quantification of colocalization was performed using the FIJI plugin under the Analyze/Cell Counter/Cell Counter tab.

For quantification of Cadherin 13 for snRNA-seq validation, we utilized Imaris. ROIs were drawn around individual granule cells manually using both DAPI and neuronal marker Neun, (image not shown) as a granule cell marker. ROIs were individually drawn around several granule cells from 3 different DG sections from 3 different animals per condition. Imaris then calculated the volume of all objects within ROIs. Values reported are the sum of Cdh 13 volume/ROI volume to normalize for the size of a cell. All ROIs are randomly taken from cells located in the granule cell layer of the dentate gyrus. Cells were selected by a different researcher and analyzed blind to treatment conditions.

For morphology experiments, Image J was used to measure rNSC length from the center of the cell soma to the longest radial process. For bushy head analysis, Imaris was used to create ROIs around individual cells that were separated enough from other rNSCs and were completely contained in the imaging section. Volumetric analysis was then performed similarly to Cadherin-13 analysis where volume within ROIs was measured. Volume data was then normalized to ROIs to control for different size ROIs.

### Optogenetic Stimulation

To stimulate cholinergic afferents from either the Medial Septum or Diagonal Band, we used optogenetics to carefully control the firing frequency and duration of activity. For neurogenesis experiments, animals were stimulated for 3 days using an 8hz paradigm. Laser intensity was set to 5 mWh and fired at 1 ms pulses for a total of 30 seconds every 5.5 minutes. There was a total of 11 five-minute stimulations per hour for 8 hours total so mice received 88 30-second stimulations during each day. This stimulation paradigm has been previously used by (Bao et al., 2017) and was shown to have circuit-specific effects for a distinct circuit projecting from the MS/DB.

Given that several studies have reported increased acetylcholine levels during the encoding/novel exploration of certain brain regions, we thought that further enhancing cholinergic activity within the DG would improve learning. Therefore, we stimulated cholinergic afferents during the encoding stage of a novel place recognition (NPR), contextual fear conditioning, or, during an open field task. For behavior experiments, animals were stimulated for a total of 10 minutes using 5mWh intensity at 1ms pulses for the entire time of the behavioral paradigm.

### Chemogenetic Stimulation

Chemogenetic activation of circuits provides the advantage of long-term circuit activation without the need to continuously stimulate. Chemogenetic stimulation was performed by one of the two following methods. For neurogenesis experiments, we administered CNO via drinking water at 2.5mg/mL for a total of 3 days. Fresh water bottles containing CNO-infused drinking water were replaced daily. For morphology experiments, we administered CNO at 1mg/kg via I.P. injection for a total of 3 days at the same time each day. Animals were perfused 1 hour after the last injection on day 3.

### Proliferation Labeling

Proliferating cells were labeled using a thymidine analog EdU similar to that described in (Quintanilla *et al*., 2019). Briefly, animals were injected I.P. with EdU at 4mg/kg 4 times on the last day of experiments. EdU was then visualized by using a click chemistry reaction which helped visualize proliferating cells in sections.

### Animal Behavior

Three different behavioral experiments were performed to test anxiety or learning/memory. All animals were injected with either AAV5-EF1a-DIO-hChR2(H134R)-eYFP virus or AAV5-EF1a-DIO-eYFP into the DB and then allowed to recover. All behavioral experiments were performed with animals attached to optical fibers to control for stimulation during the encoding phase of a task. To test anxiety animals were placed in an open field 40cmX40cmX40cm plastic square chamber and recorded while being optogenetically stimulated using the previously described paradigm for 5 minutes. Noldus Ethovision XT was used to analyze animal position during recordings. A 25cm X 25cm square was used to indicate the center of the chamber when analyzing videos. Time spent within the center and outside the center was quantified with Noldous Ethovision.

To test spatial learning, we used a Novel Place Recognition, (NPR) on the same cohort of mice used for the open field test. The NPR consisted of 5 days total with the first three days being familiarization. Animals were allowed to freely explore for a total of 5 minutes during familiarization. The square plastic chamber was cleaned between each animal using a 70% ethanol solution to mask the scent from any previously tested animal. On the fourth day of the NPR, or the encoding phase, we placed two identical objects on the same side of the chamber 20 cm apart from each other. Objects used for exploration were glass cylinders (height: 4cm, base diameter 1.5 cm) and were stuck to the chamber floor to prevent the mouse from moving them. Most animals showed no preference for any object and the location of objects was randomized across mice during the encoding and retrieval phases. Animals were allowed to interact with objects for a total of 10 minutes while being recorded and optogenetically stimulated. On the fifth/retrieval day, the position of one object was moved to the opposite side of the chamber, and animals were recorded while interacting with both objects. Videos were scored manually using a separately trained researcher who quantified the interaction time with both objects during encoding and retrieval. Any animal that did not meet the following inclusion criteria was removed from the analysis: percent exploration time < 30 % and total interaction time with both objects > 3 seconds. Percent exploration time was calculated the following way %ET = (ET_novel_ – ET_old_)/ (ET_novel_ + ET_old_). The discrimination ratio was calculated the following way DR = (ET_novel_)/(ET_novel_ + ET_old_). We also calculated the % of exploration bias by taking an absolute value of the %exploration time.

Contextual Fear conditioning was used to test contextual memory and was the last of the 3 behavioral tasks performed on each animal. For the encoding/training phase, animals were connected to optical fibers and placed in a contextual fear conditioning chamber (Med Associates) and allowed to explore for 5 minutes. Animals received two shocks that lasted a total of 2 seconds at .6mA at the 1-minute and 3-minute intervals. While animals were encoding/training they were also being optogenetically stimulated using the same stimulation paradigm described above. For each retrieval phase, animals were connected to optical fibers and allowed to explore the chamber for 3 minutes. Freezing time was assessed using a built-in camera within the chamber. Since an optical fiber was attached to animals, we used the analysis software EZtrack (Pennington et al., 2019) to assess the percent freezing time during all tasks. We used the default settings within EZtrack to analyze the freezing percentage in videos where animals had fibers. The chamber was cleaned with 70% ethanol between animals to mask the scent of the previous animal. To test the generalization of freezing, we used Context B, where we placed a white plastic cover over the floor and along the walls of the chamber, altering its appearance.

### In Vitro Electrophysiology

At 4 to 5 weeks after AAV5-EF1a-DIO-hChR2(H134R)-eYFP injections, ChAT-Cre::Nestin-GFP mice were anesthetized with isoflurane (5% in O_2_) and transcardially perfused with ice-cold aCSF (N-methyl-D-glucamine, NMDG-based solution) containing the following (in mM): 92 NMDG, 30 NaHCO3, 25 glucose, 20 HEPES, 10 MgSO4, 5 sodium ascorbate, 3 sodium pyruvate, 2.5 KCl, 2 thiourea, 1.25 NaH2PO4, and 0.5 CaCl2, equilibrated with 95% O_2_ and 5% CO_2_ (pH 7.3, 305-315 mOsm). Brains were rapidly removed, and acute coronal slices (280 μm) containing the basal forebrain or hippocampus were cut using a Leica vibratome (VT1200, Germany). Next, slices were warmed to 34.5°C for 8 minutes. Then, slices were maintained in the holding chamber containing HEPES aCSF (in mM): 92 NaCl, 30 NaHCO3, 25 glucose, 20 HEPES, 5 sodium ascorbate, 3 sodium pyruvate, 2.5 KCl, 2 thiourea, 2 MgSO4, 2 CaCl2, 1.25 NaH2PO4 (pH 7.3, 305-315 mOsm) at room temperature for at least 1 hour before recording.

Individual slices were visualized under an upright microscope with differential interference contrast (IR-DIC) video microscopy and an IR-sensitive CCD camera (Scientific, FWCAM, USA). Responses were evoked by 5-ms light flashes (473 nm) delivered through a 40X objective attached to a microscope using an LED (Thorlabs, Canada). Patch pipettes with a resistance of 4–6 MΩ were pulled from borosilicate glass capillaries (1.5 mm outer diameter, 0.86 mm internal diameter, World Precision Instruments, USA) using a micropipette puller (PC-10, Narishige, Japan). The internal solution used contained the following (in mM): 130 K-gluconate, 20 HEPES, 4 MgCl2, 4 Na-ATP, 2 NaCl, 0.5 EGTA, 0.4 Na-GTP (pH 7.2, 290 mOsm).

For circuit mapping experiments, we recorded from nestin-GFP+ cells in the dentate gyrus of the hippocampus, we recorded light-evoked currents under the voltage-clamp recordings (holding at -65 mV) by applying 8 Hz light pulses for 2 s at every 30 s. When needed, 100 μM d-(−)-2-amino-5-phosphonopentanoic acid (d-APV), 20 μM 6-cyano-7-nitroquinoxaline-2,3-dione (CNQX), and 10 μM atropine were added to block NMDA, AMPA/kainate and muscarinic ACh receptors, respectively. Light-evoked postsynaptic current amplitude was calculated at the peak of the first response after light pulses.

Recordings were conducted in the whole-cell configuration using a Multiclamp 700B amplifier (Axon Instruments, USA). Signals were filtered at 1 kHz and sampled at 10 kHz using the Digidata 1440A (Axon Instruments, USA), data acquisition was performed using pClamp 10.3 (Axon Instruments, USA). Series resistance (Rs) was monitored throughout all experiments and cells with Rs changes over 20% were discarded.

### Microdissection of Dentate Gyrus for snRNA-Seq

The Dentate Gyrus was dissected from animals that were injected with AAV5-EF1a-DIO-hChR2(H134R)-eYFP and allowed to recover. Animals were then stimulated using the stimulation protocol described previously. After stimulation, animals were transcardially perfused using ice-cold PBS solution. The brain was then removed and placed in ice-cold PBS and sliced bilaterally across the sagittal midline. Each hemisphere was then taken and the DG was dissected carefully in ice-cold PBS. To ensure that mostly stimulated DG tissue was used for sequencing experiments, only the upper half or dorsal DG was then collected. Both Dorsal DG sections from each animal were then placed in a cryogenic tube and flash frozen and placed in -80°C.

### Split-Seq Library Preparation and Sequencing

Single-nucleus RNA sequencing was performed following the SPLiT-seq method with minor modifications (Rosenberg et al., 2018). Nuclei isolated from flash-frozen DGs were performed as previously described (Qian et al., 2020; Su et al., 2017). Briefly, tissue was thawed, minced, and homogenized using a tissue grinder in a 1 mL of HB buffer (1 mM DTT, 0.15 mM spermine, 0.5 mM spermidine, EDTA-free protease inhibitor, 0.3% IGEPAL-630, 0.25 M sucrose, 25 mM MgCl2, 20 mM Tricine-KOH) for 5 to 10 strokes, then filtered through a 40 mm strainer, under layered with a cushion buffer (0.5 mM MgCl_2_, 0.5 mM DTT, EDTA-free protease inhibitor, 0.88 M sucrose) to prevent damage to nuclei, and centrifuged at 2800 g for 10 minutes in a swinging bucket centrifuge at 4°C. The pellets were resuspended in 1 mL of cold PBS-RI (1x PBS + 0.05 U/ml RNase Inhibitor). The nuclei were passed through a 40 mm strainer. 3 mL of cold 1.33% formaldehyde solution was then added to 1 mL of cells. Nuclei were fixed for 10 mins before adding 160 uL of 5% Triton X-100. We then permeabilized nuclei for 3 mins and centrifuged at 500 g for 3 mins at 4°C. Nuclei were resuspended in 500 uL of PBS-RI before adding 500 uL of cold 100 mM Tris-HCl pH 8. Then, nuclei were spun down at 500 g for 3 mins at 4°C and resuspended in 300 uL of cold 0.5 X PBS-RI. Finally, nuclei were again passed through a 40 mm strainer and then counted on a hemocytometer, diluted to 1,000,000 cells/mL. mRNA from single nuclei were tagged 3 rounds with barcoded primers, with in-cell ligations using T4 DNA ligase within 96-well plates. Plates were incubated for 30 mins at 37°C with gentle shaking (50 rpm) to allow hybridization and ligation to occur. The ligation products were purified with Dynabeads MyOne Streptavidin C1 beads. After washing beads once with 10 mM Tris and 0.1% Tween-20 solution and once with water, beads were resuspended into a solution containing 110 uL of 2X Kapa HiFi HotStart Master Mix, 8.8 uL of 10 mM stocks of primers BC_0062 and BC_0108, and 92.4 uL of water. PCR thermocycling was performed as follows: 95°C for 3 mins, then five cycles at 98°C for 20 s, 65°C for 45 s, 72°C for 3 mins. After these five cycles, Dynabeads beads were removed from the PCR solution and EvaGreen dye was added at a 1X concentration. Samples were again placed in a qPCR machine with the following thermocycling conditions: 95°C for 3 mins, cycling at 98°C for 20 s, 65°C for 20 s, and then 72°C for 3 mins, followed by a single 5 mins at 72°C after cycling. Once the qPCR signal began to plateau, reactions were removed.

PCR reactions were purified using a 0.8X ratio of KAPA Pure Beads and cDNA concentration was measured using a qubit. For tagmentation, a Nextera XT Library Prep Kit was used. 600 pg of purified cDNA was diluted in water to a total volume of 5 uL.10 uL of Nextera TD buffer and 5 uL of Amplicon Tagment enzyme were added to bring the total volume to 20 uL. After mixing by pipetting, the solution was incubated at 55°C for 5 mins. A volume of 5 uL of neutralization buffer was added and the solution was mixed before incubation at room temperature for another 5 mins. PCR was then performed with the following cycling conditions: 95°C for 30 s, followed by 12 cycles of 95°C for 10 s, 55°C for 30 s, 72°C for 30 s, and 72°C for 5 mins after the 12 cycles. 40 uL of this PCR reaction was removed and purified with a 0.7X ratio of SPRI beads to generate an Illumina-compatible sequencing library.

### Single Nuclei RNA-seq data analysis

FASTQ files were deconvoluted and transcript abundance was estimated using zUMIs v2.9.6 (Parekh et al., 2018) and GENCODE vM26 annotations (Harrow et al., 2012). Bases 1-66 were extracted from the R1 file, corresponding to cDNA sequence, and for the R2 file, bases 1-10 represented UMI, and bases 11-18, 49-56, and 87-94 represented the three barcodes. Internally, zUMIs performs a two-pass alignment using STAR v2.7.3a (Dobin et al., 2013) with the additional parameter “--limitSjdbInsertNsj 2000000”. Barcodes and UMIs were also filtered for quality using “num_bases=1, phred=10”. We then generated cell x genes matricies. Data were then imported into R v3.6 and Seurat V3 (Stuart et al., 2019) objects were created for each individual sample before being merged. Cells were filtered such that there were at least 1000 UMIs and 500 genes detected, and fewer than 10% of the transcripts were mitochondrially contributed. Data were then scaled and normalized using scTransform (Hafemeister and Satija, 2019) and integrated using Seurat. Clusters were identified using Louvain-Jaccard clustering with multilevel refinement (resolution=2) performed on the top 100 PCs. Marker genes of each cluster were identified using the FindMarkers function in Seurat, and differential expression analysis was performed using RNA CPMs in R using Wilcoxon Rank-Sum test. Differentially expressed genes were considered significant with a nominal p-value < 0.01.

We determined whether DB cholinergic afferent activation regulated DG cells by intersecting our data with gene ontology databases using g:Profiler2 (Kolberg et al., 2020) and looking at GO terms associated with Biological Processes (BP) only.

### Inference of developmental trajectories

Pseudotime trajectory of analysis was performed with Slingshot (Street *et al*., 2018), filtering out lowly expressed genes and using a 2-dimensional embedding based on PCA of Z-scores of gene expression patterns in RGL (green), RGL-Like (blue), and Neuroblast (red) cells in control (darker hues) and stimulated (lighter hues). We identified 1 lineage that gives rise to Granule cells for our RGL cluster.

### Niche Net Analysis

To identify active ligands and their regulatory effects on to rNSCs, we used Niche Net. Niche Net analysis was performed following the established pipeline in (Browaeys *et al*., 2020). Briefly, we used our merged Seurat object used to identify cell clusters and DEG’s. Our Seurat object was then subset based on the cell types being analyzed using Seurat V4 (Hao et al., 2021). Analysis was performed by selecting our rNSC cluster as the receiver, and the remainder of the niche as the senders, excluding CA clusters and subiculum clusters. Cell classes were used to analyze our data instead of clusters to reduce the number of comparisons required for analysis and increase the generalization across cell classes. Top ligand and receptor pairs were identified by using our existing DEGs in each cluster. The same DEG threshold, p-value < .01, and test, Wilcoxon Rank-Sum test, was used during Niche Net analysis.

## Data Availability

Single-nucleus RNA-seq data have been deposited to the Gene Expression Omnibus (GEO) under accession GSE202481. The repository will be released publicly upon acceptance, but reviewers can access it using the token (Provided after publication).

## Code Availability

Code is available in a Song Lab Repo on GitHub at https://github.com/lquin003/Song-Lab_Quintanilla_et_al.

## Statistical Methods

All statistics were performed using R version 4.1, Seurat Version 3.1 or 4, or PRISM 9. Individual animals are treated as biological replicates unless otherwise noted in figure legends. Data are all presented as mean +/- S.E.M. unless indicated in figure legends. Sample sizes were determined based on the number of animals used in previous publications. All manual data analysis was performed blinded to experimental genotypes including quantification of neural stem cells and behavioral analysis. Certain experiments were not possible to blind such as the quantification of anterograde traces, or bushy head and soma projections. Significance was defined as *p < 0.05, **p < 0.01, ***p < 0.001, ****p < 0.0001. No data were excluded unless specifically specified.

---

**Supplemental Figure 1:**
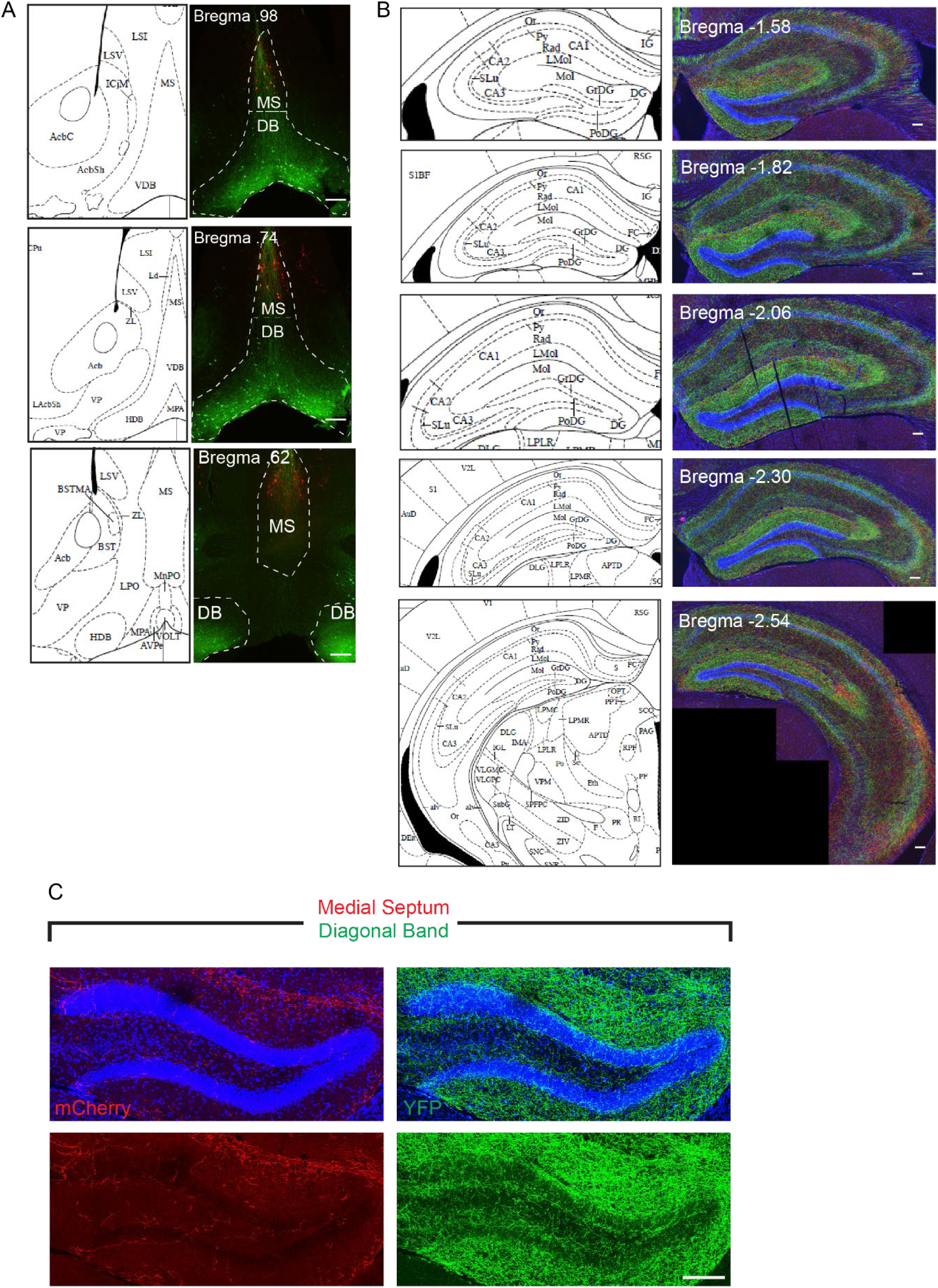
Distinct patterns of cholinergic projections from Medial Septum and Diagonal Band of Broca to DG. A) Representative confocal image of injection site in animals injected with 2 color fluorophores. MS-mCherry, DB-YFP. Scale bar = 100 µM B) Representative confocal images of hippocampal sections used to quantify volumetric projections spanning about 2 mm of the hippocampus. Scale bar = 100 µM C) Representative confocal image showing close up of cholinergic projections from the MS or DB to the DG. Scale bar = 100 µM

**Supplemental Figure 2:**
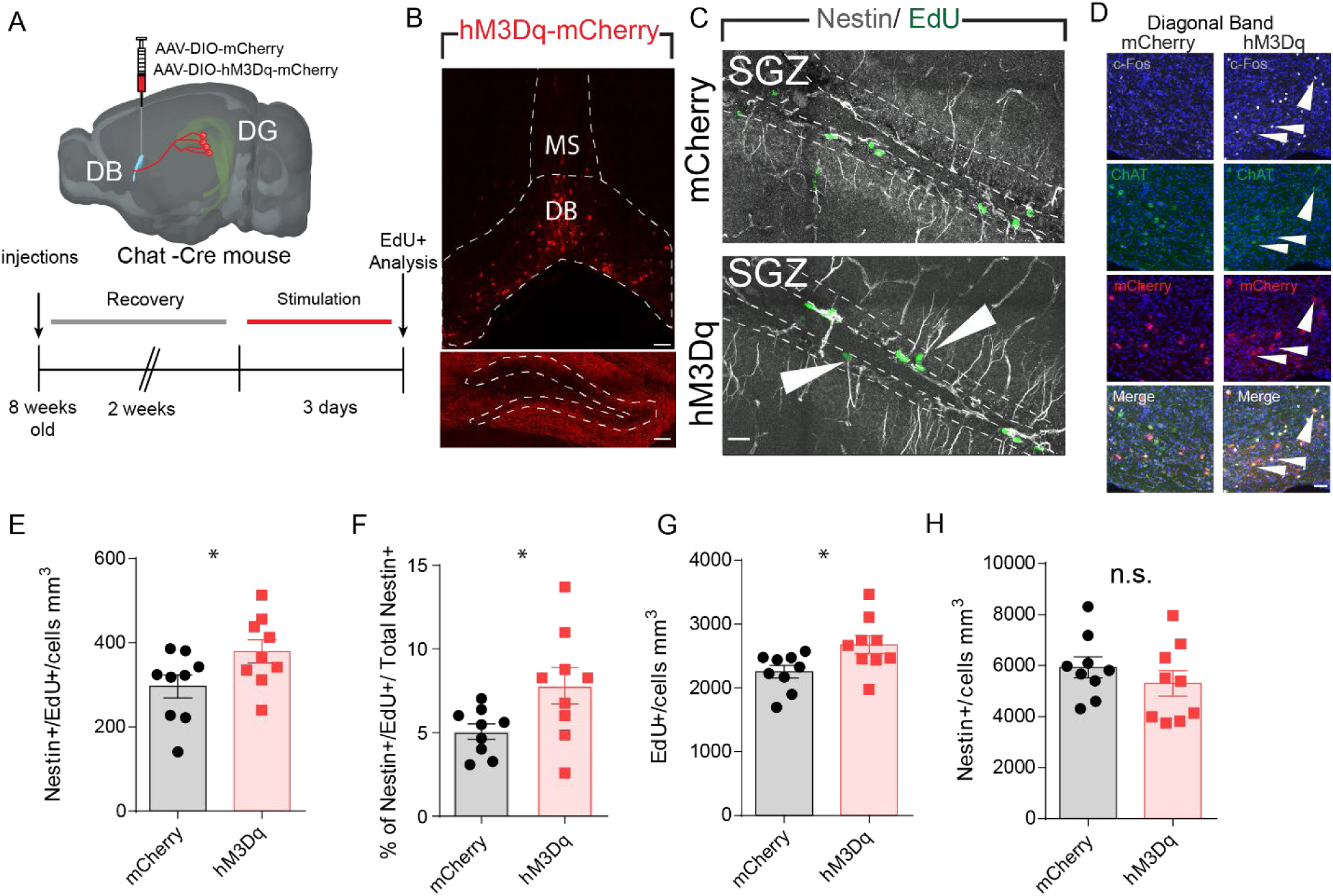
Chemogenetic activation of DB neurons activates rNSCs. A) Schematic of timeline and experimental setup for chemogenetic stimulation of DB cholinergic afferents and rNSC proliferation analysis. B) Immunofluorescence image of DB targeted AAV viral injection and hM3Dq afferents in the DG. DB scale bar =100µm, DG Scale bar = 100 µm. C) Representative immunofluorescence section of SGZ used to quantify rNSC proliferation with Nestin/EdU. Scale bar = 50 µm. D) Representative immunofluorescence image of chemogenetically activated DB cells stained with ChAT, mCherry, and c-Fos antibodies. Scale bar = 50 µm E) Quantification of proliferating neural stem cells. Bars indicate mean +/- S.E.M. n= (9,9), p-val =.0497 by student’s t-test . F) Quantification of percent of proliferating neural stem cells. Bars indicate mean +/- S.E.M. n= (9,9), p-val =.0348 student’s t-test. G) Quantification of overall proliferation in the SGZ. Bars indicate mean +/- S.E.M. n = (9,9), p- val =.0266 by student’s t-test. H) Quantification of neural stem cell pool. Bars indicate mean +/- S.E.M n = (9,9).

**Supplemental Figure 3:**
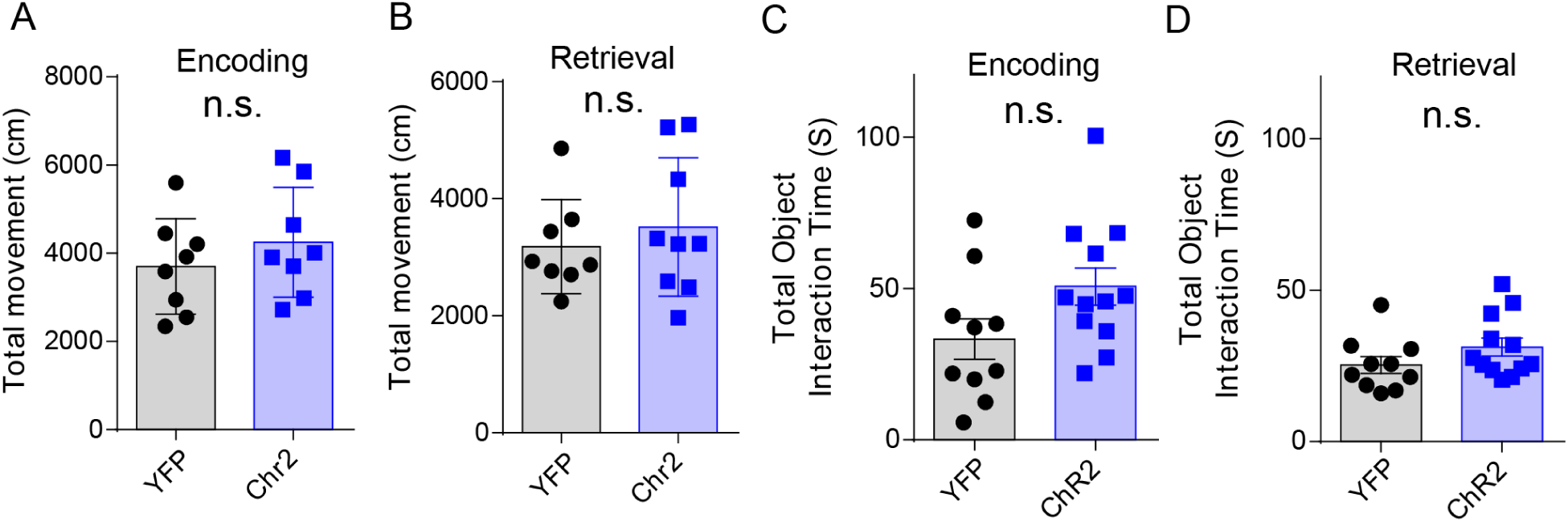
Stimulating DB-DG projections does not affect locomotion. A) Quantification of total movement during encoding or retrieval (B). Bars indicate mean +/- S.E.M. n = 8 C) Quantification of total time interacted with both objects during encoding or retrieval (D). Bars indicate mean +/- S.E.M. n = (10,12)

**Supplemental Figure 4:**
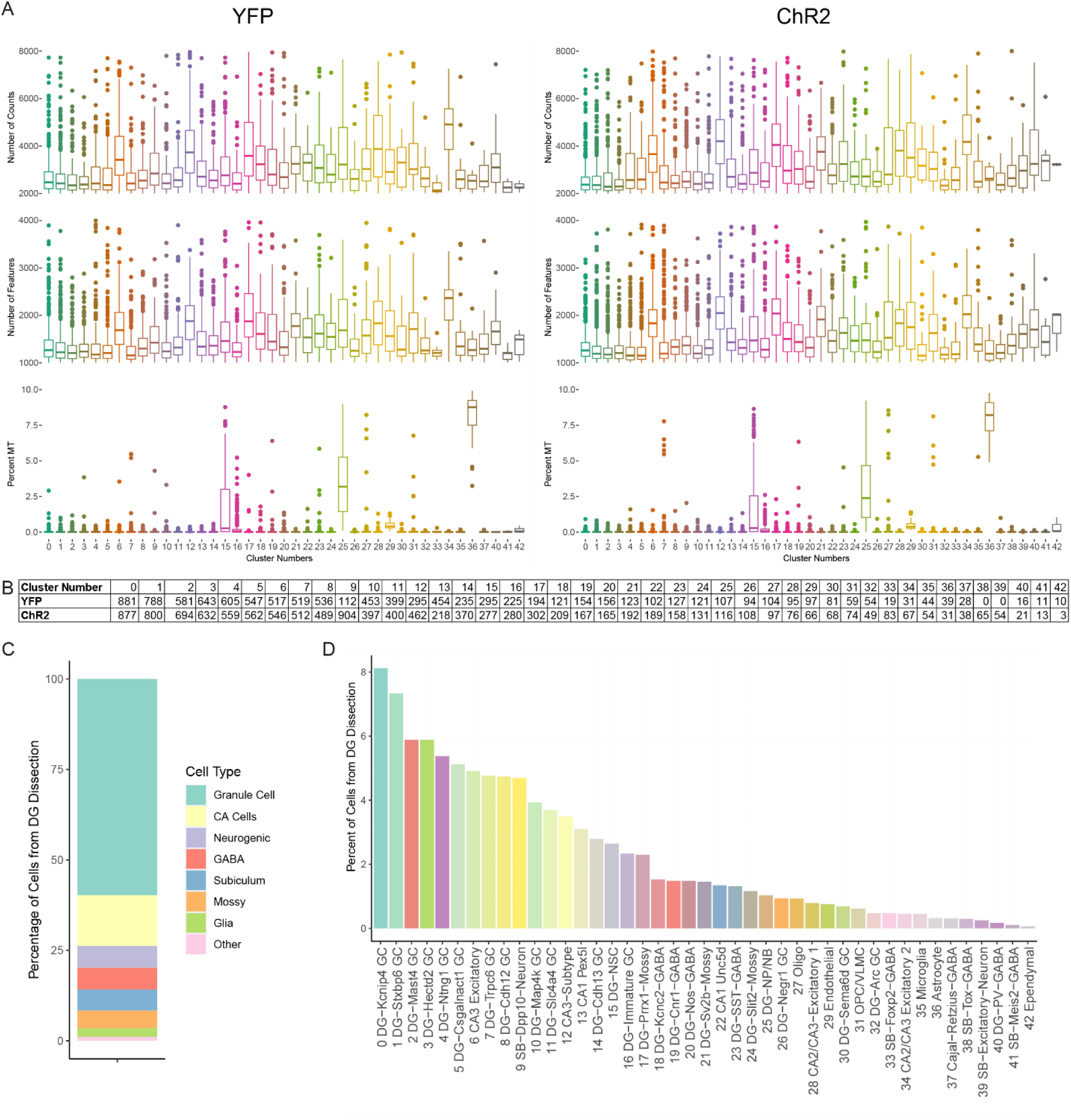
Quality control of snRNA-seq clusters. A) Boxplots showing upper quartile, median, and lower quartile of the total number of counts (UMIs), number of features (genes), or percent of mitochondrial reads for each cluster separated by YFP and ChR2. Cells that did not meet the minimum requirements for any of these 3 criteria were excluded. B) Total number of cells in each cluster for both Control and ChR2 that passed quality control metrics. C) Percentage of cell classes for all clusters dissected from the DG. D) Percentage of cells within each cluster from all dissected cells from our experiment.

**Supplemental Figure 5:**
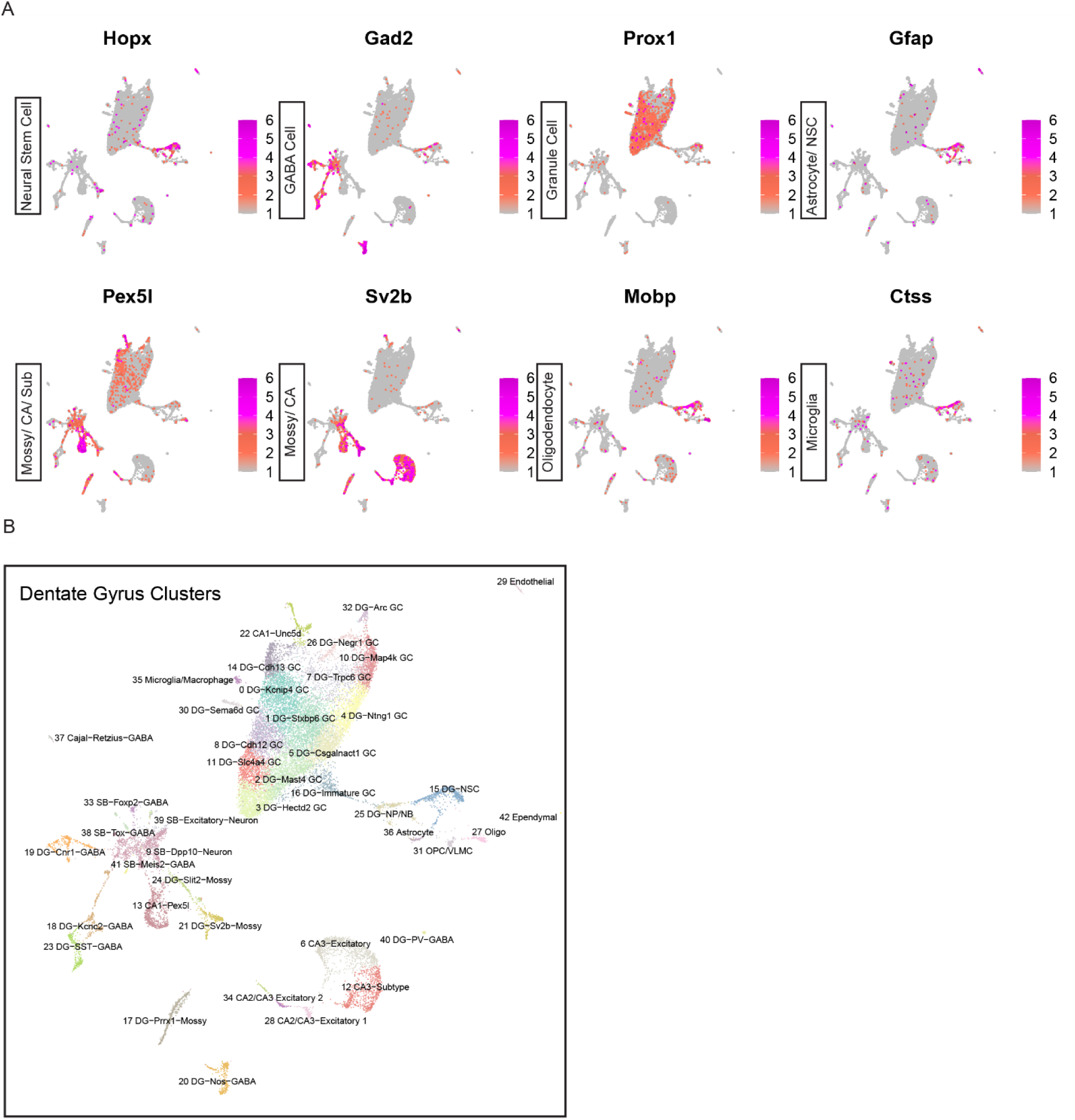
Establishment of cluster identities using marker genes. A) UMAP plots showing expression of different cell-type-specific markers. Plot title shows labeled marker gene and legend shows expression. B) Dimension reduction analysis of our 42 DG clusters identified from snRNA-seq. Cluster number and location are labeled above corresponding clusters.

**Supplemental Figure 6:**
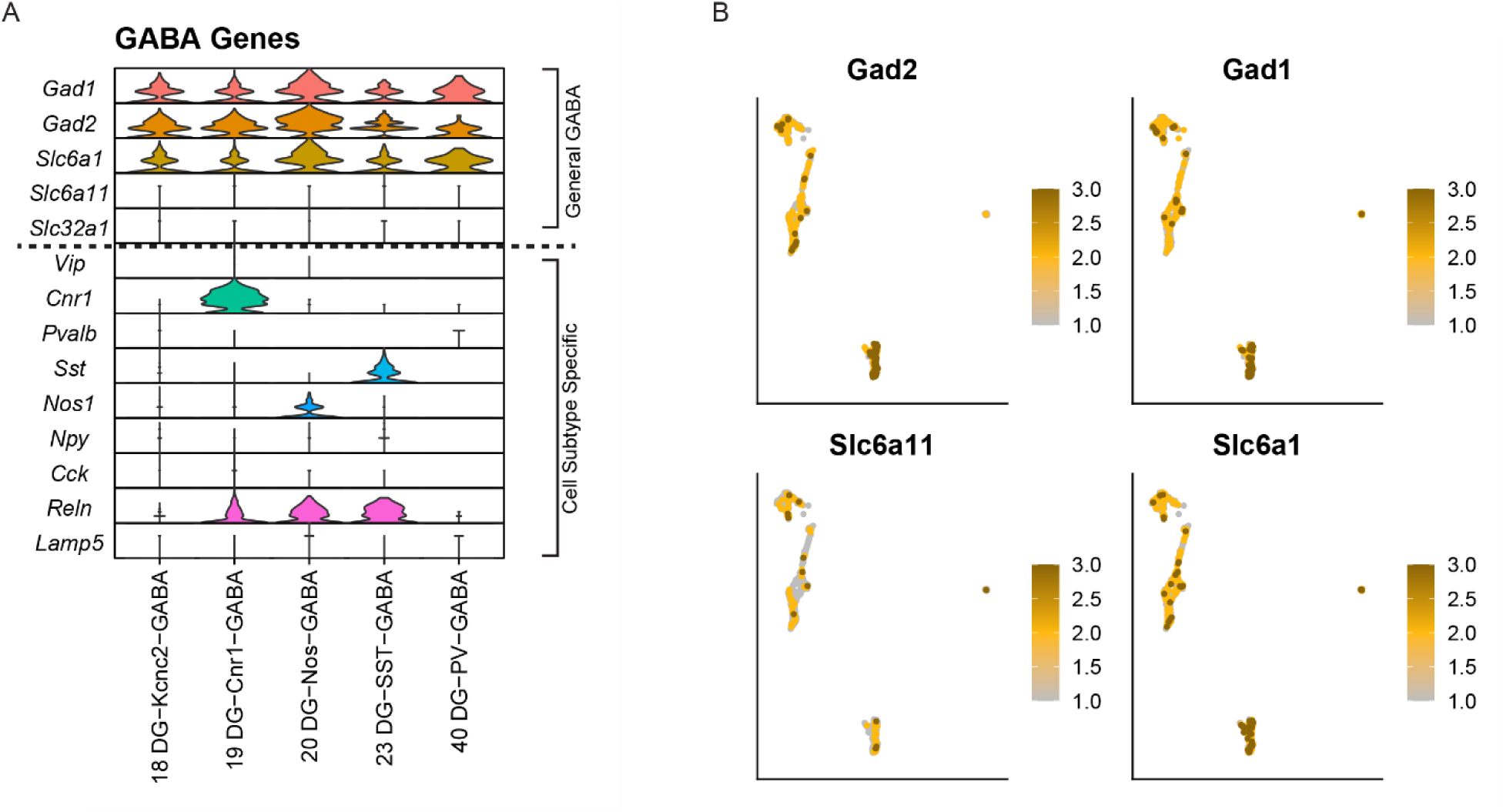
Identification of GABA interneuron subclasses. A) Violin plot showing major GABA markers that are both shared and exclusive to certain cell types in clusters. Marker genes below indicate cell-type-specific markers. B) UMAP plots showing expression of 4 general GABA markers in DG-GABA clusters subset from the original clusters.

**Supplemental Figure 7:**
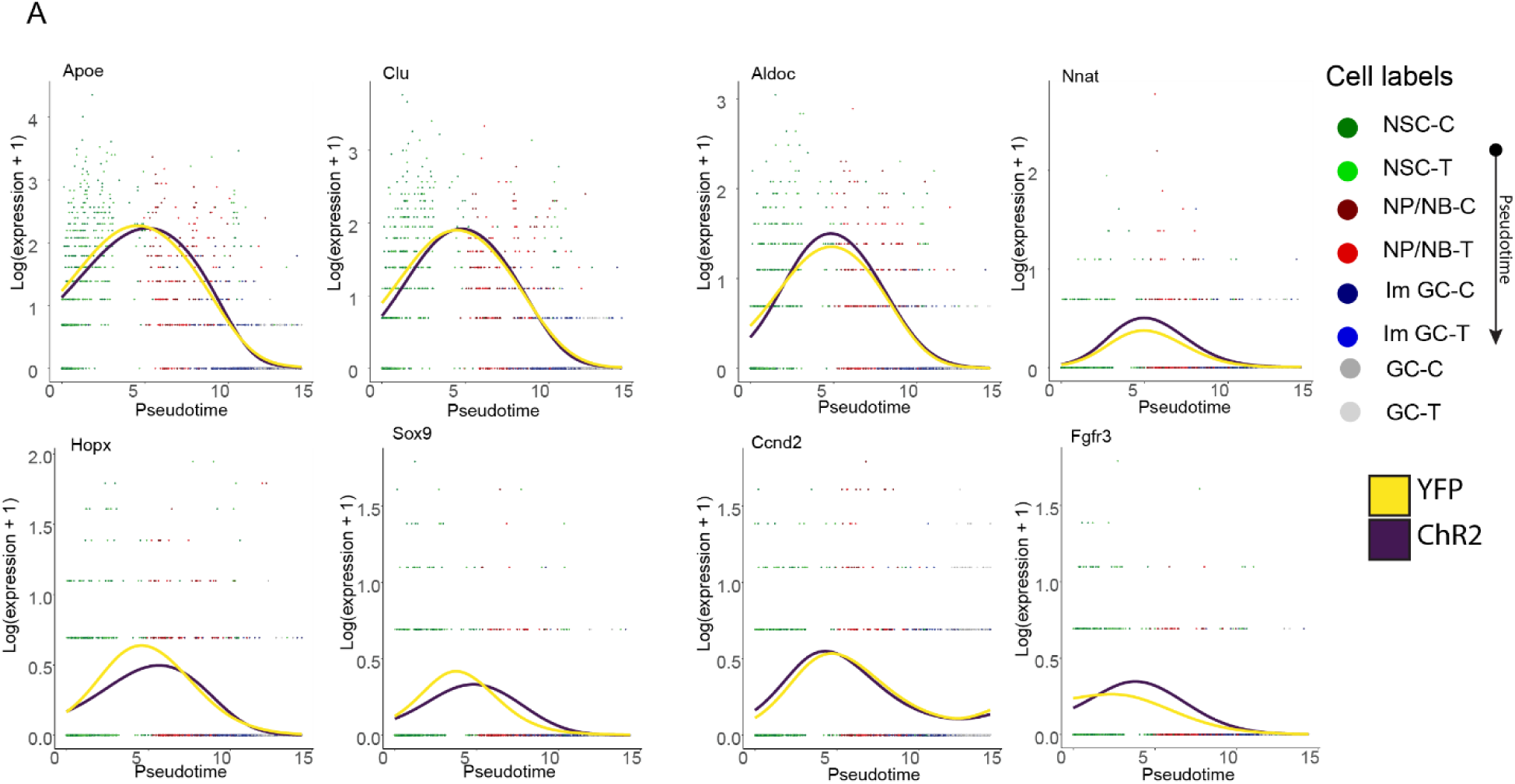
Pseudotime analysis of neurogenic lineage. A) Pseudotime representation of neurogenic clusters generated using Slingshot. Pseudotime goes from left to right. YFP (control) clusters are darker and ChR2 (treatment) clusters are lighter.

**Supplemental Figure 8:**
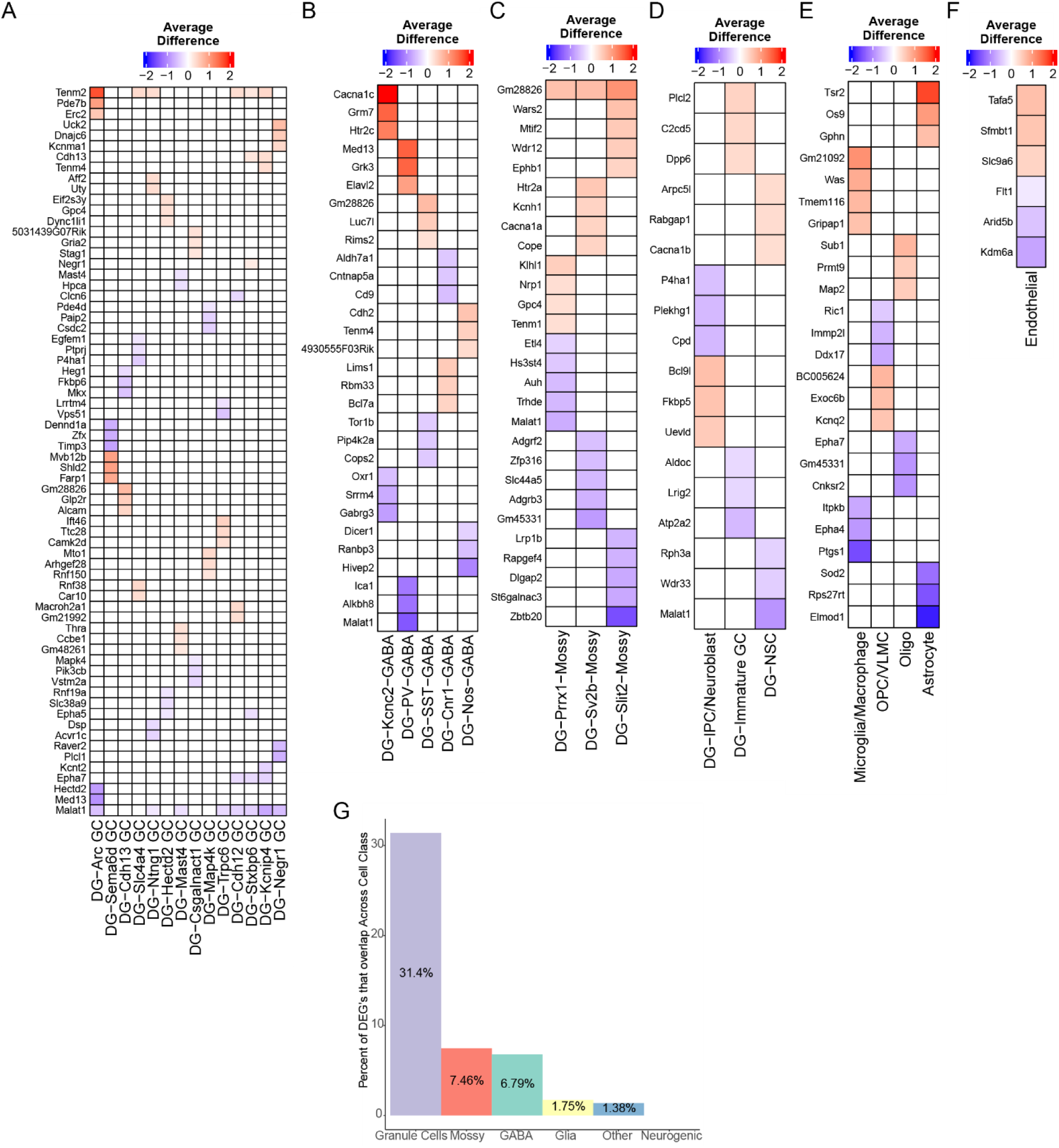
Visualizing the most significant DEGs in each cluster. A) Heatmap plot showing the average difference of most significant DEG’s in each direction in each cluster grouped in the Granule cell class, B) GABA cell class, C) Mossy cell class D) Neurogenic cell class E) Glia cell class F) Other cell class G) Percent of overlapping genes in at least 2 clusters across cell classes.

**Supplemental Figure 9:**
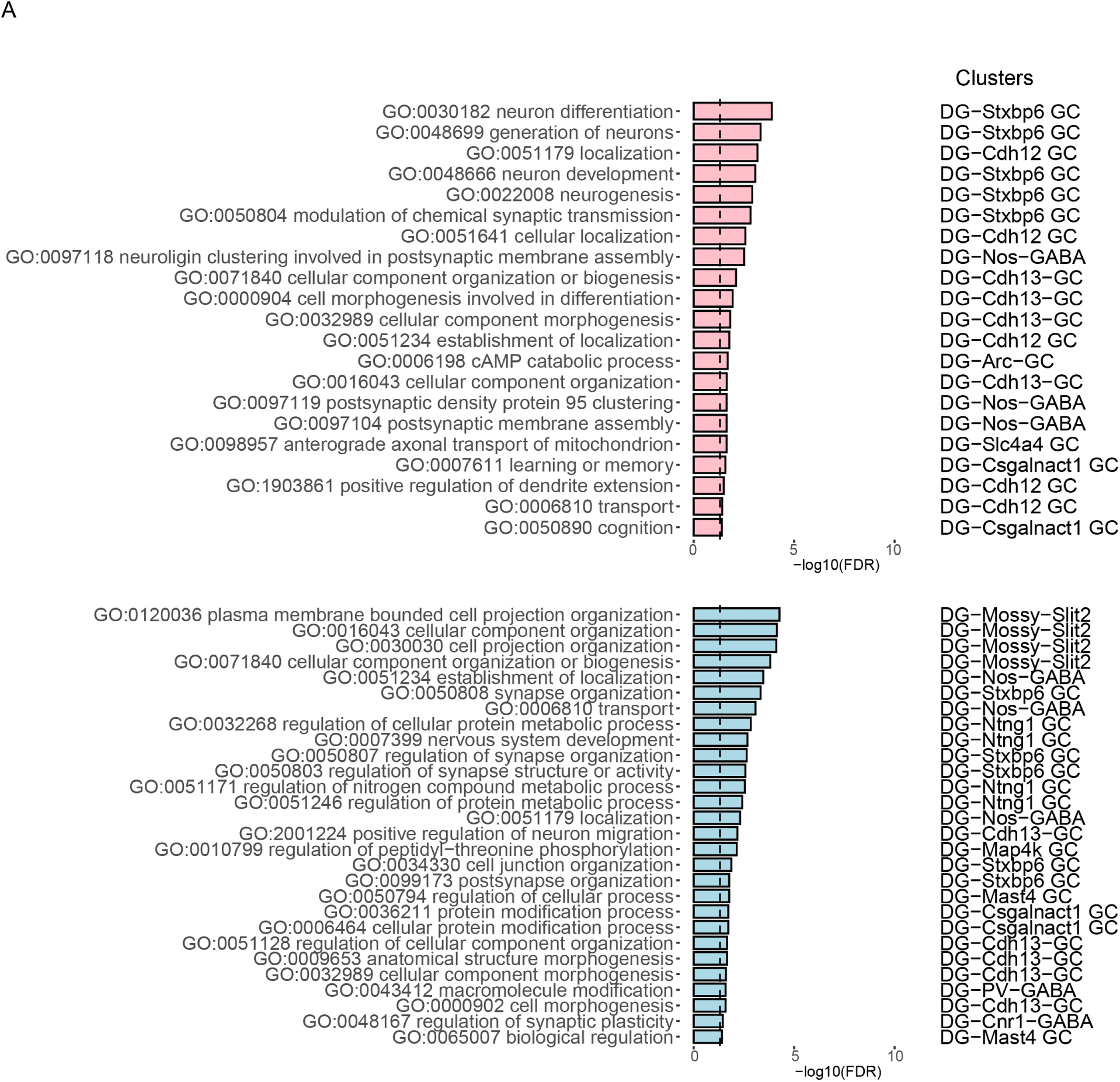
Top-ranked GO terms for neuronal clusters not in Figure 6. A) Bar plots showing the most significant GO terms for remaining neuronal clusters not shown in Figure 6. GO terms are ranked from most significant to least significant and split between upregulated and downregulated DEGs from each cluster.

**Supplemental Figure 10:**
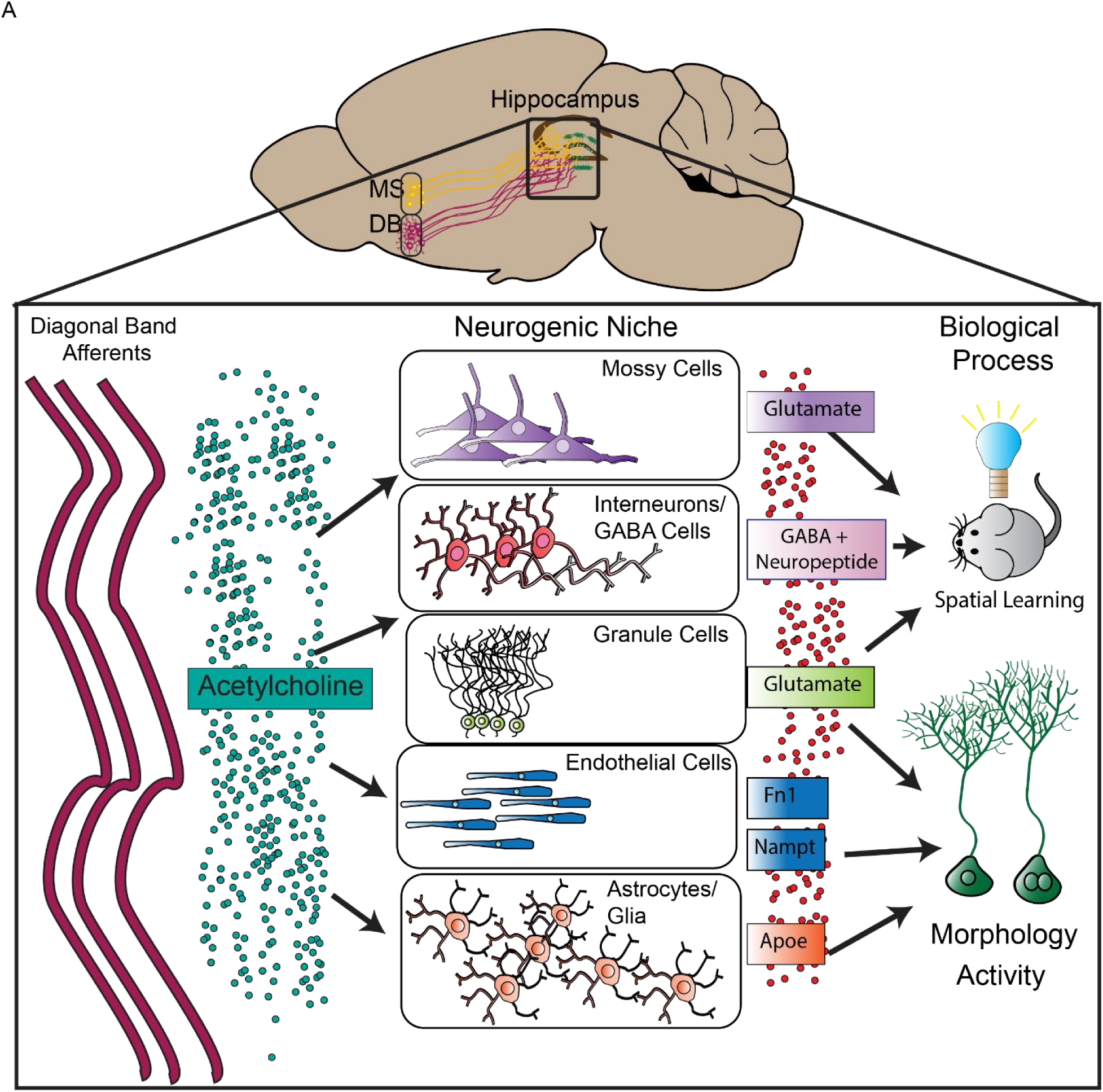
A working model for cholinergic circuit-dependent regulation of distinct hippocampal functions. A) Schematic of working model summarizing all the findings from this study supporting both spatial learning and rNSC morphology and proliferation.

## Notes

https://github.com/lquin003/Song-Lab_Quintanilla_et_al.

### Competing Interest Statement

The authors have declared no competing interest.

